# Computational Framework for Identifying Ion Channel Mutation-Compensating Interventions

**DOI:** 10.64898/2026.09.20.753002

**Authors:** Hananel Hazan, Michael Levin

**Author notes:** **Competing Interest Statement**: The Levin Lab receives support via a Sponsored Research Agreement with Morphoceuticals, a company seeking to commercialize bioelectric repair strategies.

## Abstract

We present an automated pipeline for finding therapeutic interventions in channelopathies. It starts from patch clamp recordings of the variant channel, searches over the pharmacologically accessible conductances, and says which currents must change and by how much. The intervention never touches the mutated channel: it compensates by modulating others. The feasibility of this approach is demonstrated through two computational strategies, each addressing a specific capability gap. First, we employ high-fidelity NEURON simulations combined with exploratory search algorithms to model two clinically described variants, and identify “stability windows” in which channel modulations restore the reference firing pattern. The compensating configurations are degenerate: 37 conductance triples fire the same number of action potentials at every one of the 35 injected-current levels, and the 10,780 triples reaching the best intervention value produce only six distinct firing patterns. Second, to address the computational cost of such detailed modeling, we implement a differentiable Hodgkin-Huxley model in PyTorch. Applied here to theoretical mutation screening, this approach trades granular detail for speed, treating the search as a continuous optimization problem that makes very large, non-local searches of the conductance parameter space affordable: the differentiable forward model evaluates 34,000–64,000 candidate parameter sets per second on a single consumer graphics card, against 11.9–16.9 per second for the non-differentiable pipeline on a 72-core node. The gain arises from forward-model throughput feeding a search rather than from the gradient itself, and it is what makes the 32,000,000-point survey of the solution topology reported here affordable. These computational predictions can inform practical drug development and high-throughput screening by naming the currents that must change and by how much.

## 1 Introduction

Ion channelopathies, disorders caused by mutations in voltage-gated ion channels, represent a significant challenge in neurological therapeutics [1–15]. Alongside genetic causes, numerous syndromes targeting nervous system function are caused by environmental or pre-natal exposure to a wide variety of chemical agents, stress, and other factors [16–27]. Ion channel defects, at the genetic, transcriptional or functional level, are not limited to the nervous system. The emerging field of developmental bioelectricity [28–32] has identified numerous contexts in which ion channel malfunction induces structural birth defects [3, 33–37], limits healing and regeneration potential [38–40], is involved in aging and senescence [41], and potentiates cancer [1, 42–50]. Other examples in which electrophysiological dynamics play an important role in health and disease include inner ear, gut, kidney, intestine, retina, immune response, and numerous other body systems [51, 52]. Bioelectric status is therefore an attractive point of intervention across these systems [51, 53].

Despite advances in understanding these mechanisms [31, 54–56], developing effective treatments remains complex due to the intricate relationship between channel mutations, physiological dynamics, and the resulting cell- and circuit-level function. This is complicated by the fact that bioelectric regulation offers complexity and emergent dynamics at two levels. First, individual channels can open and close in place (as a result of numerous regulatory signals). Reading their mRNA or even protein expression level, or regulating their expression, is often not sufficient to gain predictive control of cellular voltage state. Second, ion channels form circuits which have collective behaviors such as compensating for one another or forming feedback loops (as in voltage-sensitive channels, which are in effect transistors). In many tissues, such as the brain and central nervous system, these properties are essential, and they underline complex computation necessary for adaptive function. In disease states, the same dynamics make treatment difficult.

Gene therapy and similar approaches are limited because they target the cellular hardware, which is important but does not encompass the “software” aspects of the complex electrical dynamics that run on electrically active media. This is in addition to the known difficulties with gene therapy in general [57, 58]. Thus, there is great need for approaches that induce compensatory changes at the level of electrophysiology. Specifically, rather than targeting the mutated channel itself, which may be resistant to repair, we propose a compensatory strategy: using pharmacological agents to modulate the conductance of other, non-mutated channels. By tuning these healthy channels, the neuronal circuit can be brought back into a functional range. This is known to be possible due to the inherent nature of the plasticity of electrophysiological networks. For example, recent work in animal models in vivo has shown that methods to modulate bioelectric state can improve wound healing [59–62], rescue chemical and genetically-induced birth defects [63, 64], induce regeneration of appendages [38–40], and exert other forms of control over growth and form in the context of chemical, genetic, or mechanical damage [65, 66].

Pharmacological agents available (and often already human-approved) to target ion channels, pumps, and gap junctions form a powerful toolkit of electroceuticals for a wide range of indications [67–69]. However, traditional drug discovery approaches often struggle to account for the dynamic nature of ion channel and circuit behavior. The existing successes in identifying powerful triggers of complex downstream states via computational methods [70–78] need to be expanded and enabled to exploit powerful emerging methods of AI.

Here, we introduce a dual-strategy framework that bridges the gap between biophysical understanding and therapeutic intervention. We present two distinct computational methods, each addressing a specific capability gap in drug discovery:

1. **High-Fidelity Simulation (NEURON):** This method utilizes the NEURON simulator [79] to incorporate granular, patient-specific ion channel interactions. While it offers the highest biological realism necessary for personalized medicine, the simulation is non-differentiable and computationally intensive, requiring resource-heavy search algorithms to identify interventions.
2. **Differentiable Simulation (Gradient Descent):** To overcome the computational bottlenecks of exhaustive search, we present a second approach using a differentiable Hodgkin-Huxley model. We recreated the original Hodgkin-Huxley equations in a differentiable environment using PyTorch [80, 81], which enables parameter tuning through gradient-based optimization. While it trades some granular detail for speed, its differentiable nature allows for rapid gradient-based optimization. We demonstrate that this method can evaluate candidate parameter sets at a rate between three and four orders of magnitude above the non-differentiable pipeline’s production rate, on the same unit of work (Section 3.7), bringing high-throughput screening within reach.

By combining published patch clamp recordings of the variant channels with advanced computational modeling, our approach offers a systematic method for identifying potential therapeutic targets. As illustrated in Figure 1, our methodology proceeds from patient cell samples through electrophysiological characterization to computational analysis and drug target identification. We parameterize our models from these recordings, which measure ion channel behavior in both healthy and affected cells.

**Figure 1:**
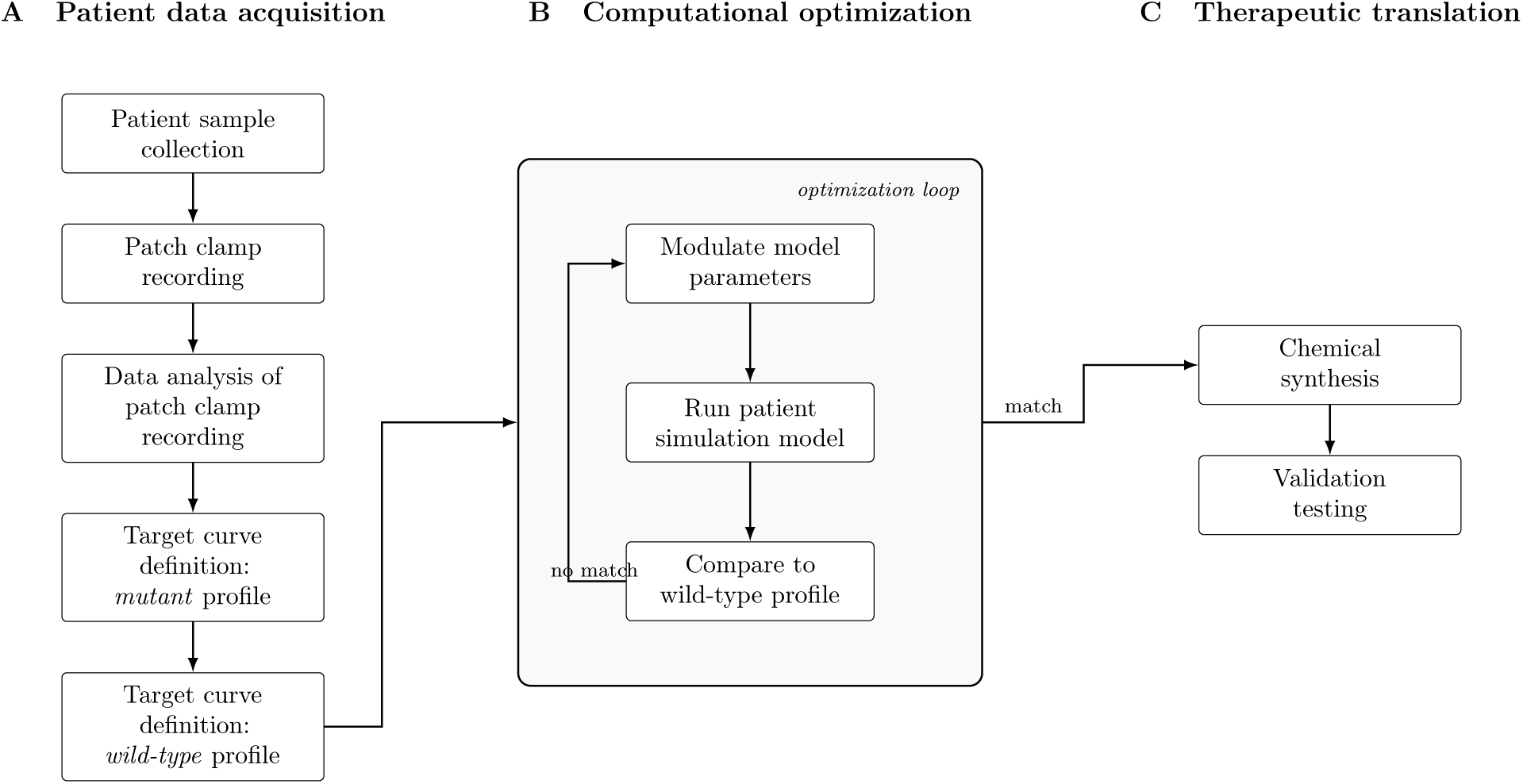
Integrated workflow for automated discovery of bioelectric interventions. **(A)** Cells from a patient carrying the channelopathy are characterised by patch clamp, and the recordings define two target curves: the uncompensated variant profile, against which the effect of an intervention is measured, and the reference profile the intervention aims to restore. **(B)** The optimization loop modulates the conductances of pharmacologically accessible channels (never the mutated channel), simulates the resulting cell, and compares it against the reference, iterating until the comparison is satisfied. The loop is run by gradient descent or by heuristic search, and the two can in principle be combined. **(C)** The resulting parameter set names the currents that must change and the magnitude of modulation required, which is the starting point for compound selection or synthesis and subsequent validation *in vitro*.

Previous research has primarily focused on electrophysiological characterization of ion channels or mutant cells [82, 83], computational models of cell-level or circuit-level disorders [84], and drug screening for novel ion channel-targeting compounds [85]. Integrating these efforts into a single flow remains an open need [86]. Our goal here is a methodology in which patch clamp data taken from patient samples feeds directly into a computational search for interventions, without separate modeling phases and without losing the patient-specific information along the way. The pipeline combines these approaches so that candidate interventions can be evaluated systematically [82, 83] while accounting for biological variability, which is also what makes it a starting point for personalized therapeutic approaches to ion channel disorders.

The pipeline is not specific to a single mutation or channel family: it requires only a conductance-parameterised model and a reference behaviour to restore, a condition met by the non-neural bioelectric contexts described above as well as by the neuronal ones tested here, and it carries a computational prediction through to a prescription that names the currents to change and by how much. Here we apply it to two clinical channelopathy variants in both the high-fidelity and the differentiable model, and find that the configurations restoring reference behaviour form extended sets: 37 conductance triples reproduce the variant’s own firing pattern exactly, and stability windows exist in which multiple therapeutic combinations achieve the same functional restoration, which is what makes them reachable by an intervention.

## 2 Methods

### 2.1 Overview of the Hybrid Computational Strategy

Identifying therapeutic interventions for channelopathies is limited by the cost of the forward simulation. We therefore use two models with complementary properties. High-fidelity NEURON models supply the biological realism needed to judge a candidate intervention, and carry the two case studies in Sections 3.1, 3.2, 3.4 and 3.5 and the multi-compartment results of Section 3.6. A differentiable Hodgkin-Huxley model implemented in PyTorch trades biological detail for throughput, evaluating candidates in batch on a graphics processor; it is characterised in Section 3.7 and is what makes the 32,000,000-point topology survey of Section 3.3 affordable. The two are used separately here; Section 4.2 sets out how they would be combined, with the fast model narrowing the region that a high-fidelity search then resolves. In the investigations that follow, the high-fidelity and differentiable components each identify compensating conductance changes for a specific neuronal mutation, the differentiable one by search over its fast forward model.

### 2.2 Differentiable Simulation Framework

To enable rapid optimization, we implemented a differentiable Hodgkin-Huxley model using the PyTorch [80, 81] framework, in the same class as other differentiable biophysical simulators [87]. Unlike standard simulators, this implementation allows for the calculation of gradients via automatic differentiation. We defined a loss function based on the Mean Squared Error (MSE) between the mutant and wild-type voltage traces. Optimization was performed using the Adam optimizer, allowing for the direct update of conductance parameters (*ḡ*_Na_, *ḡ*_K_, *ḡ*_leak_) via gradient descent to minimize the loss function. The implementation is tested under PyTorch 1.9 through 2.12, and every rate reported in Table 1 was measured under PyTorch 2.10.0 built against CUDA 12.8. Learning rates were selected per optimizer from a documented pre-sweep, taking for each the scale that gave the lowest squared-error loss after 120 iterations; the selected potassium rates span 0.003 to 30, and each is an interior point of the swept grid, so no optimizer is evaluated at the edge of its range.

**Table 1:** Batching and kernel fusion, not the accelerator alone, account for the differentiable model’s throughput. One candidate parameter set is one conductance triple simulated across all 35 injected-current levels, 300 ms each, at a 0.01 ms integration step. Every figure is a measured rate on the hardware named; none is projected. Graphics-card rates are medians of five launches, with the range across launches beside each.

| Pathway | Hardware | Implementation | Sets/s | Range, 5 runs |
| --- | --- | --- | --- | --- |
| Non-differentiable (NEURON) | one processor core | per-candidate loop | 1.0 | — |
| Non-differentiable (NEURON) | 72-core node | spike-count stage | 11.9 | — |
| Non-differentiable (NEURON) | 72-core node | time-warping stage | 16.9 | — |
| Differentiable | one processor core | unbatched, 1 set at a time | 0.089 | — |
| Differentiable | one processor core | batched, 1,024 sets | 20.8 | — |
| Differentiable | RTX 2070 (2018) | batched, unfused | 543 | 542.8–543.0 |
| Differentiable | RTX 2070 (2018) | batched, graph-captured | 572 | 571.7–571.7 |
| Differentiable | RTX 2070 (2018) | batched, fused | 34,279 | 34,275–34,288 |
| Differentiable | RTX 4070 Ti (2023) | batched, unfused | 2,053 | 2,046–2,057 |
| Differentiable | RTX 4070 Ti (2023) | batched, graph-captured | 2,806 | 2,790–2,809 |
| Differentiable | RTX 4070 Ti (2023) | batched, fused | 63,649 | 63,633–63,651 |
| Differentiable | RTX 4070 Ti (2023) | batched, fused, cache-resident | 100,443 | 98,001–104,926 |

*The seven graphics-card rates are medians of five independent end-to-end launches of the same benchmark on the hardware named, with the full range across the five in the last column; the five processor-core rates are single measurements and carry no range. The single-core NEURON figure is rounded down from a measured 1.10 sets per second. Four qualifications bound the comparison and are set out in Section 3.7.*

#### Algorithm 1

The guards on lines 4, 7 and 8 keep one diverged configuration from affecting the rest of its batch. One update of the conductances on the differentiable model, as used for the batched fits behind the loss-measure, optimizer and learning-rate comparisons reported here. Line 7 recovers a negative conductance and line 8 catches what the clamp leaves, since a comparison against a non-finite value does not evaluate true. Line 4 sanitises only the value passed to back-propagation, so line 3 still records the non-finite loss

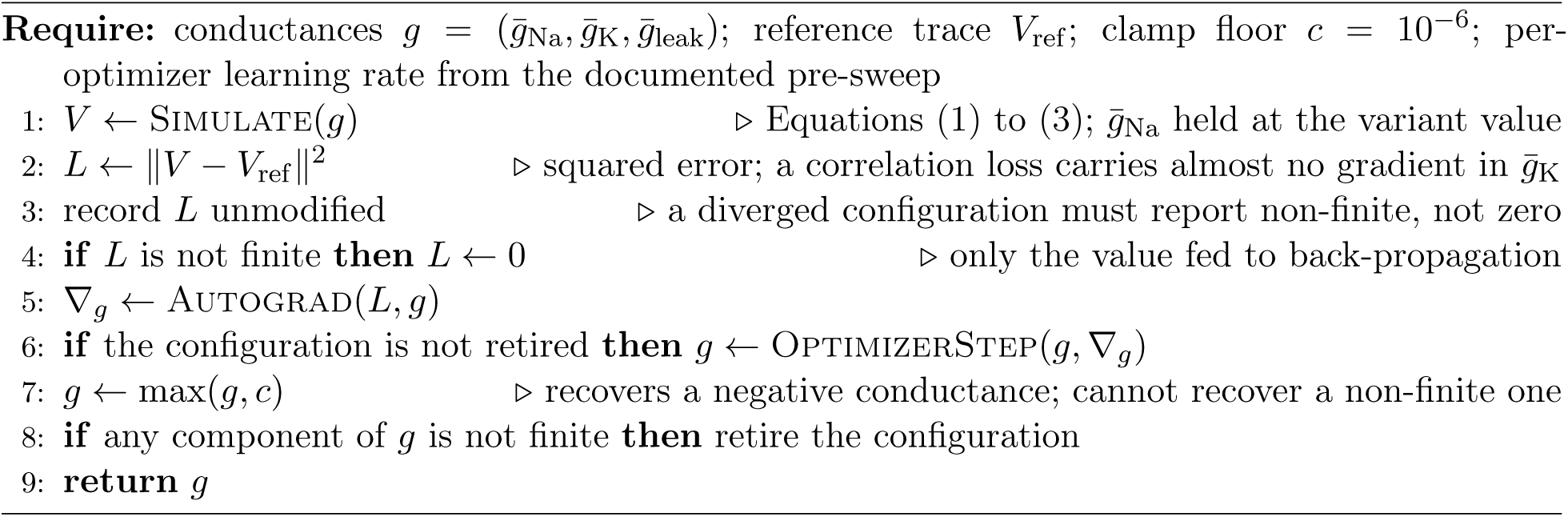

The leak rate is fixed at 0.02 times the potassium rate throughout, the ratio used in the reference implementation. The optimizer runs hold the sodium conductance at the variant value and start from a common point, and are therefore deterministic and carry no seed; the stochastic components of this study, the sampling-scheme comparison and the genetic-algorithm search, are seeded per repeat at the budgets reported with each.

The choice of loss measure determines which conductances the gradient is able to move. A correlation-based loss is invariant to affine rescaling of the simulated trace, which is principally what the potassium conductance does, and it therefore carries almost no gradient in that direction: at the start point the partial derivative of a Pearson-correlation loss with respect to *ḡ*_K_ is smaller than that of the squared-error loss by roughly four orders of magnitude (*−*0.0016 against *−*19.7), and under the correlation loss the potassium conductance remains essentially unchanged while the other parameters converge normally. We therefore use a squared-error measure, and recommend it wherever the optimized parameters include conductances whose principal effect is on trace amplitude.

Three implementation details are necessary for reproduction. The rate functions *α_n_* and *α_m_* each take the indeterminate form 0*/*0 at *V* = *−*50 mV and *V* = *−*35 mV respectively, and a simulated trajectory passes arbitrarily close to both in normal operation; we evaluate each by its series expansion within a small threshold of those potentials, with limiting values 0.1 and 1.0 ms*^−^*^1^. Separately, conductances are clamped after each update to a small positive floor of 10*^−^*^6^; without it, seven of the eight optimizers we tested, all but Adadelta, drove a conductance non-positive. An explicit finiteness test follows the clamp, since a comparison against a non-finite value does not evaluate true, and a configuration that fails the test is retired rather than stepped again. Finally, when configurations are fitted in a batch, a non-finite loss is replaced by zero before back-propagation, which keeps one diverged configuration from affecting the gradients of the others sharing its batch; the recorded loss is the unmodified one, so a divergence is reported rather than scored as zero. Algorithm 1 gives the order in which the guards apply.

### 2.3 Model Development and Simulation Framework

We developed and validated our computational approach using two distinct neuronal models derived from patch-clamp recordings. The first model examined a novel R859C mutation in the Nav1.1 sodium channel, identified in a four-generation family with generalized epilepsy with febrile seizures plus (GEFS+) [88]. The second model investigated a Kv7.2 (KCNQ2) variant causing benign familial neonatal seizures, using the published multi-compartment CA1 pyramidal neuron model of Miceli et al. [89] (ModelDB accession 118986), in which a single mechanism carries the M-current and is supplied in wild-type and D212G variant forms.

### 2.4 Parameter Analysis Framework

Our analysis focused on three key parameters: sodium channel conductance (*ḡ*_Na_), potassium channel conductance (*ḡ*_K_), and passive leak channel conductance (*ḡ*_leak_). These parameters were selected based on their fundamental role in the Hodgkin-Huxley equations. The simulation framework employed two measurement methods to assess membrane voltage quality: spike count analysis and dynamic time warping (DTW) [90]. Each intervention was evaluated against a battery of input currents to assess neuronal response patterns.

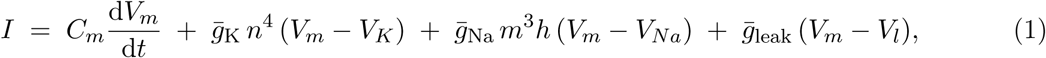

Equation (1) is the Hodgkin-Huxley membrane equation used for the differentiable simulation. It expresses the total membrane current *I* as the sum of a capacitive term, set by the membrane capacitance *C_m_*, and the ionic currents carried by the sodium, potassium and leak conductances. A mutation is represented in this pipeline as a fixed pathological alteration of these kinetic parameters or conductances, for example a reduced *ḡ*_Na_; a drug intervention is represented as a scalar multiplier applied to the maximal conductance of a channel that is not mutated, *ḡ*_K_ or *ḡ*_leak_. Solving the equation over many such multipliers is what identifies the conductance values that restore a reference voltage trajectory.

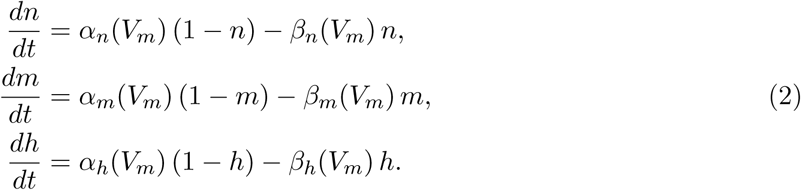

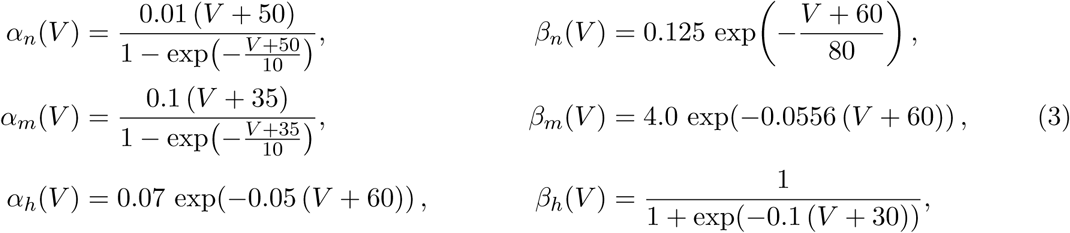

Equation (2) gives the state transitions of the gating variables *n*, *m* and *h*, which set the fraction of channels open at any instant, and Equation (3) the voltage-dependent rate functions behind them. The rate coefficients *α*(*V_m_*) and *β*(*V_m_*) are non-linear functions of membrane potential that govern the speed of transition between open and closed states. A channelopathy is modelled by altering specific coefficients within these equations. The R859C variant, for example, is represented by altering the voltage dependence of the *α_m_* activation coefficient in both midpoint and slope, so as to approximate the measured activation relationship of the variant channel: in the implementation used here, a +6.1 mV shift in the half-activation midpoint together with an approximately 34% change in the slope factor. The 8.5 mV figure reported in the source literature is a midpoint shift alone, which a pure translation of the curve does not reproduce. Because these rate functions are continuous, the partial derivative of the loss with respect to a conductance *g* can be propagated through the non-linear dynamics to the voltage trace, which is what makes gradient-based optimization of the conductances possible.

### 2.5 Intervention Strategy Development

The archived R859C exploration used a genetic algorithm for initial broad exploration, followed by uniform random sampling over the parameter box and retention of a fixed number of best-ranked candidates. Six genetic-algorithm configurations were launched, with populations of 1,000, 10,000 and 20,000 and generation limits of 10,000 and 100,000, all using a crossover probability of 0.5, a mutation probability of 0.3 or 0.5, Gaussian mutation and tournament selection. One completed: 10,000 generations at a population of 1,000, using 6,002,343 evaluations. The others stopped between 10 and 8,989 generations. This stage was exploratory and its results do not enter the analysis reported here, which rests on the uniform random sampling described next. Each conductance was drawn from a range centred multiplicatively on its baseline value: a factor of 50 either side for the leak conductance, and a factor of 5,000 either side for the two gated conductances. For the R859C variant this produced 38,850 candidate interventions, formed by pairing each of 1,050 mutant-type to wild-type (MT-WT) candidates (difference vectors from a variant configuration to a reference one) with each of 37 mutant-type to mutant-type (MT-MT) configurations, the alternative parameter sets that reproduce the observed variant behaviour. Each difference vector is applied to each variant configuration in both directions, added and subtracted. A result with a negative conductance is rejected; a conductance of exactly zero is admitted.

The Kv7.2 search, built for this study, draws candidates log-uniformly. Because the box is defined multiplicatively and spans several orders of magnitude, drawing uniformly in the logarithm of the conductance samples each factor-of-ten interval with equal density. Figure 2 compares the two schemes on a common budget of 1,000,000 draws, with eight independent repeats per condition. Log-uniform draws located a compensating configuration in every repeat, at a median first hit between 2,905 and 58,245 draws depending on the variant and a slowest single repeat of 231,941; uniform draws located one in no repeat within that budget. Two properties of the log-uniform draws account for that difference. They place just over a quarter of candidates within a factor of ten of the baseline sodium conductance, where the compensating configurations lie, against about one draw in five hundred under uniform sampling; and they raise the share of draws that produce a spiking cell from 5.7% to 38.2% for the sodium variant and from 17.0% to 42.4% for the potassium variant. The two act separately: for the larger potassium shift the schemes are within 1.4 percentage points of each other on the spiking share, 46.8% against 45.3%, and the reach in that condition follows the decades the draws occupy.

**Figure 2:**
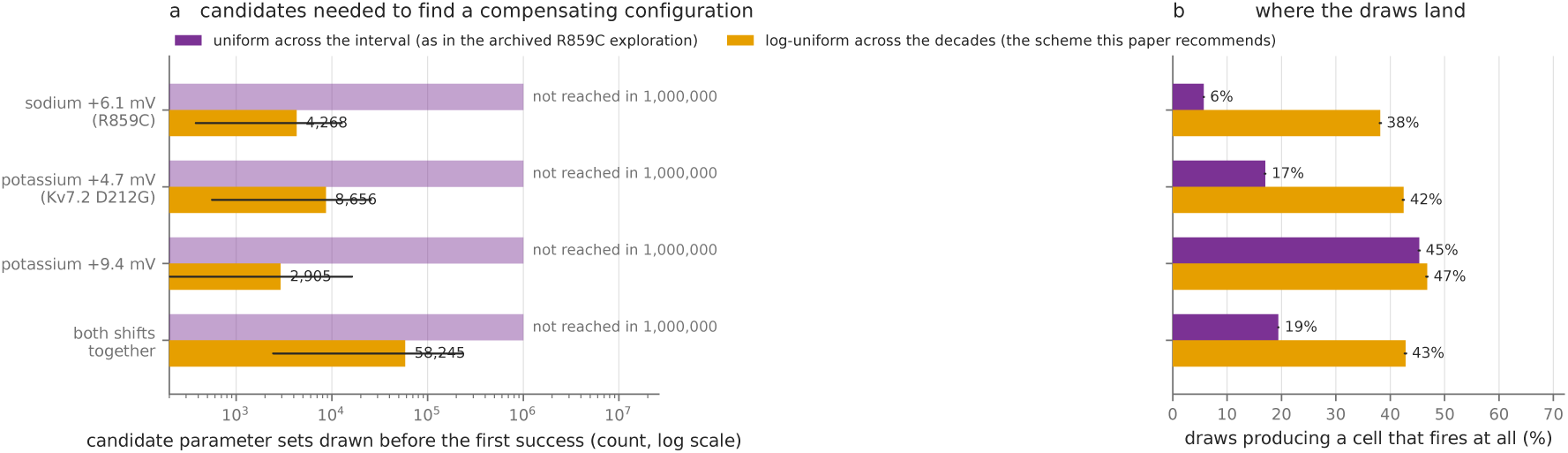
Sampling the conductance box log-uniformly reaches compensating configurations that uniform sampling does not. The box is defined multiplicatively and spans several orders of magnitude, so the way it is sampled decides what a finite budget can reach; this comparison is the evidence for the log-uniform scheme recommended here. Both schemes were run on the differentiable model over the pipeline’s own parameter box, with eight independent repeats per condition, each with its own seed and a budget of 1,000,000 draws, and success defined as reaching within 5 percentage points of the best value known for that variant, scored on spike count across all 35 injected-current levels. **(a)** Candidates drawn before the first success, logarithmic axis; the bar is the median across repeats and the horizontal line the full spread. Uniform draws did not locate a compensating configuration in any of the 32 repeats, so the hatched bars are drawn at the budget and are lower bounds rather than measurements; log-uniform draws succeeded in all 32, with medians from 2,905 to 58,245 draws and a slowest single repeat of 231,941. **(b)** The share of draws that produce a spiking cell. For the larger potassium shift the two schemes are within 1.4 percentage points of each other on this measure, so in that condition the reach is carried by which decades the draws occupy.

To assess similarity between wild-type and mutated neurons under parameter modifications, we employed two complementary metrics. Our primary measure was the MSE between the spike counts of the two cells across the injected-current ladder (Figure 3). We supplemented it with the DTW distance (Figure 4), which captures temporal patterns in neuronal activity that spike counting alone does not resolve.

**Figure 3:**
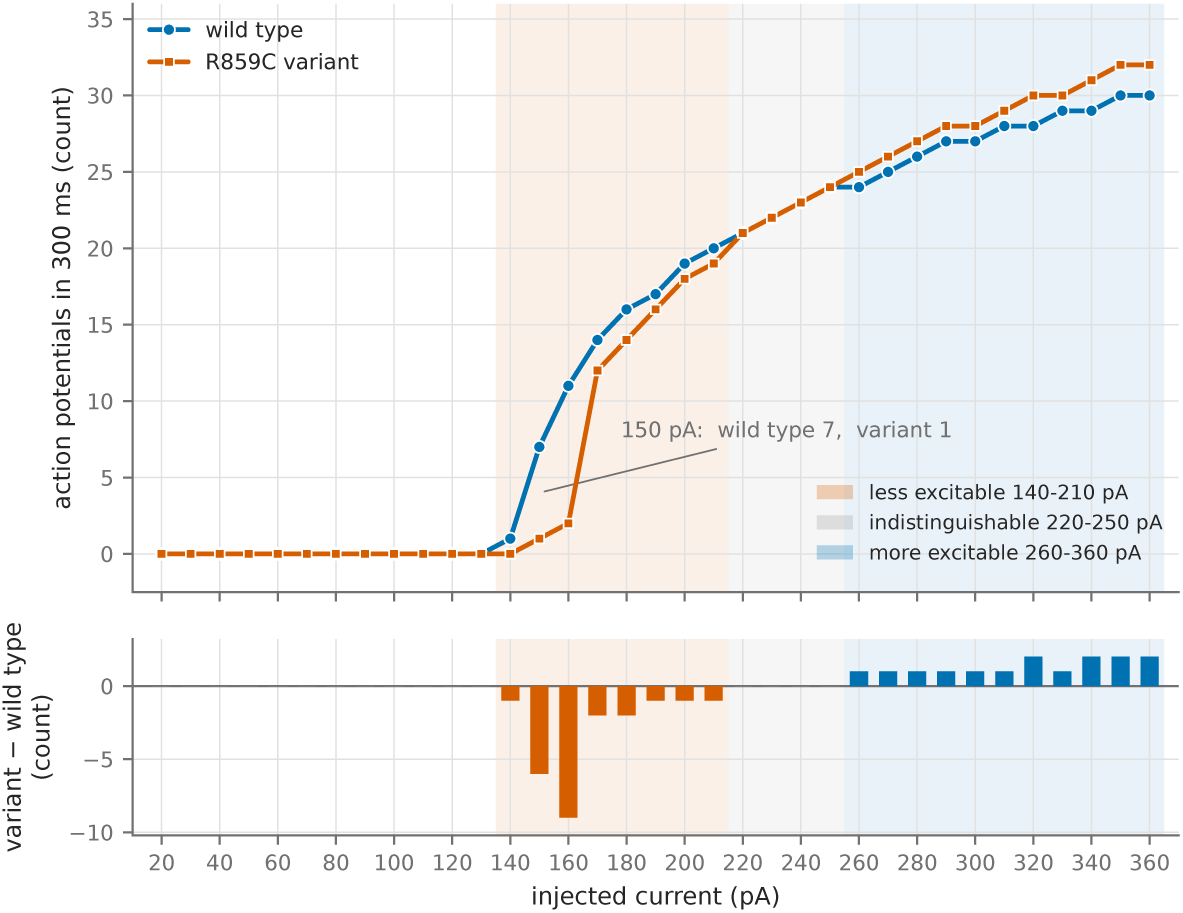
The R859C variant shifts the threshold for repetitive firing rather than abolishing the capacity for it. Action potentials counted in the 300 ms sweep for the reference and variant cells at each of the 35 injected-current levels, 20 to 360 pA in 10 pA increments, under somatic current clamp with a 50 ms delay and a 200 ms step; spikes were counted with the analysis pipeline’s own detector. **Upper panel:** spike count against injected current, with the three regions shaded. **Lower panel:** the per-level difference, variant minus reference. The deficit is regional: fewer action potentials from 140 to 210 pA, an identical number at every level from 220 to 250 pA, and more at all eleven levels from 260 to 360 pA. Summed over the ladder, the reference fires 498 action potentials against the variant’s 490. The whole ladder is shown because a regional threshold shift and a global loss of excitability are different therapeutic targets.

**Figure 4:**
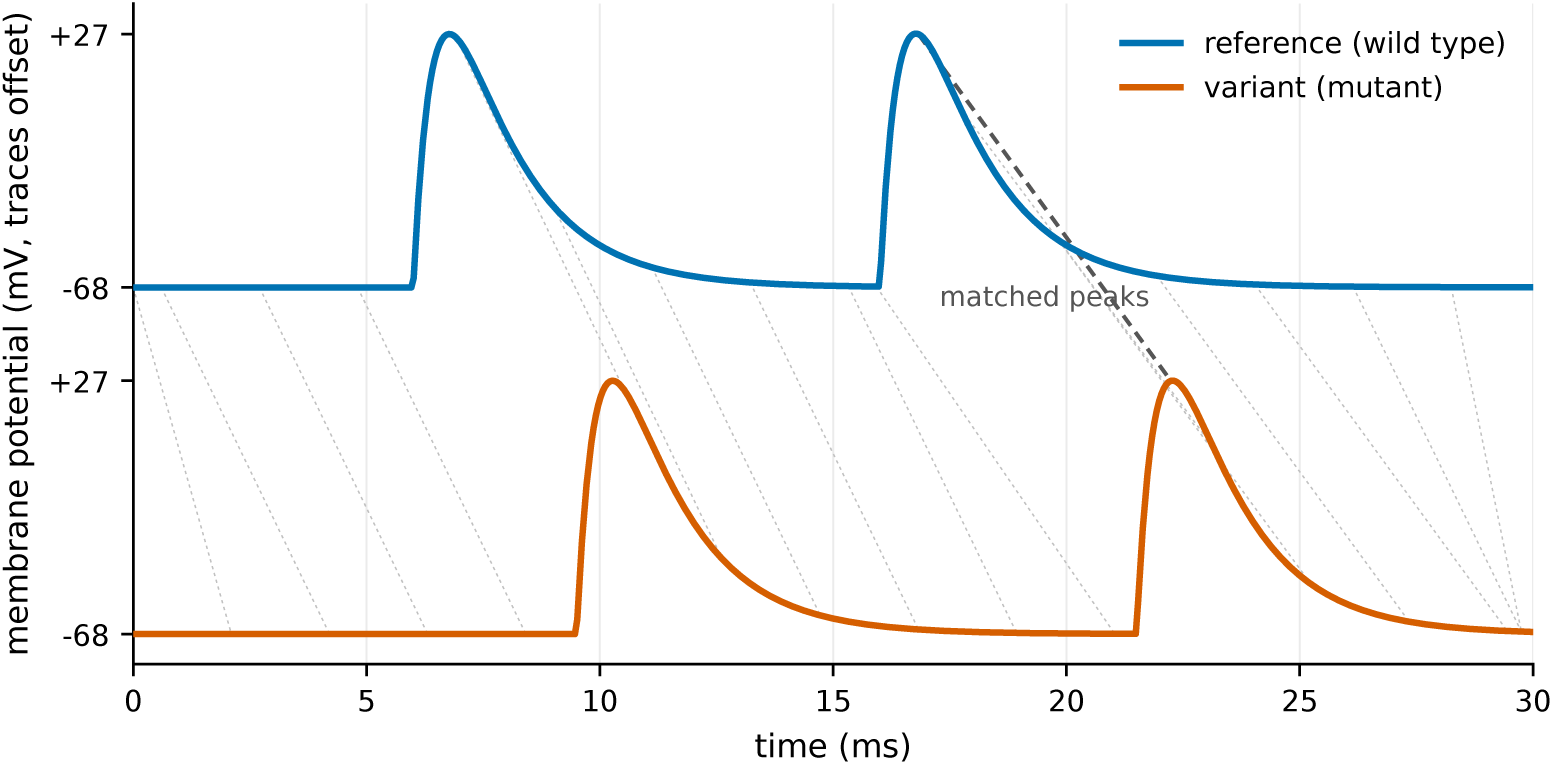
Dynamic time warping separates a cell that fires the right pattern late from one that fires the wrong pattern. Spike count measures how *often* a cell fires but not *when* or in what shape; dynamic time warping supplies the missing comparison by aligning two traces non-linearly in time before measuring the distance between them. Grey lines show the warping path, computed between the two traces shown rather than drawn schematically; the heavier dashed line marks the correspondence between matched action-potential peaks. The two traces are offset vertically for legibility.

The DTW distance addresses the oversimplification inherent in a spike-count MSE. It compares the voltage traces themselves, level by level across the injected-current ladder, and allows for stretching and shifting in time, which gives a more detailed picture of the similarity between the wild-type trace and the trace of the treated mutant cell.

Throughout, similarity is reported as a normalized percentage against the untreated variant:

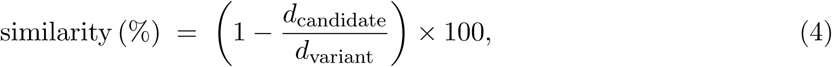

where *d*_candidate_ is the distance between the treated cell and the reference under the chosen measure and *d*_variant_ is the untreated variant’s own distance from the reference, with the scale floored at *−*100%. On this scale 0% is the untreated variant and 100% is behaviour indistinguishable from the reference; negative values denote a candidate worse than no treatment, and the floor at *−*100% is reached when a candidate leaves the cell twice as far from the reference as no treatment. For the R859C study under the spike-count measure the denominator *d*_variant_ is 37 action potentials, being the summed absolute difference in spike count between reference and variant across all 35 injected-current levels, measured against the reference the search integrated alongside every candidate. This quantity is distinct from the 37 MT-MT configurations described above, with which it should not be confused. The reference it is measured against is one of three that appear in this work, each with its own scale; Section 3.5 names all three.

The intervention framework was constructed by calculating the differential between selected MT-MT and MT-WT candidates, providing precise conductance adjustments required for phenotype correction. This approach enabled systematic evaluation of intervention efficacy across varied neuronal states while accounting for biological variability. Intervention success was quantified on a normalized scale against wild-type neuron behavior, with careful attention to stability characteristics and reproducibility of conductance modifications.

## 3 Results

Our computational framework for identifying ion channel mutation interventions follows a systematic progression from characterizing mutant phenotypes to developing targeted therapeutic strategies. We validated this approach through three complementary investigations that collectively demonstrate the feasibility of automated drug discovery for channelopathies.

The first investigation focuses on detailed characterization of a specific mutation’s electrophysiological impact, using the well-documented R859C mutation in Nav1.1 channels associated with generalized epilepsy with febrile seizures plus (GEFS+). Building upon this characterization, our second investigation expands the scope to parameter space exploration, examining how systematic conductance modifications can potentially restore normal neuronal function in benign familial neonatal seizures. The final investigation presents a differentiable forward model whose throughput makes large non-local searches of the conductance space affordable, and asks what gradient descent on that model adds. The three investigations run from mutation characterization, through targeted intervention discovery, to a general-purpose search tool.

### 3.1 The R859C Variant Reduces Sodium Channel Availability and Shifts the Threshold for Repetitive Firing

To characterize the functional impact of the R859C mutation [88] and establish baseline parameters for intervention discovery, we simulated the electrophysiological properties of this mutation, implementing the previously characterized model [88] identified in a four-generation family with generalized epilepsy with febrile seizures plus (GEFS+). The R859C mutation neutralizes a positively charged arginine in the domain 2 S4 voltage sensor of the Nav1.1 channel subunit, a residue conserved across mammalian sodium channels and lower organisms, suggesting critical functional importance.

We simulated wild-type and R859C mutant neurons using the NEURON simulator, implementing the published model from ModelDB (model #87585) based on the Barela et al. characterization [88]. The heuristic search evaluated 38,850 intervention candidates across the injected-current ladder, pairing each of the 1,050 MT-WT candidates with each of the 37 MT-MT configurations. Each simulation tested neuronal responses using both spike count analysis for quantitative assessment and the DTW distance to evaluate membrane voltage quality and temporal dynamics. The stimulus protocol was somatic current clamp with a 50 ms delay, a 200 ms step and 300 ms total duration at a 0.01 ms integration step, across 35 levels from 20 pA in 10 pA increments.

Barela and colleagues reported three electrophysiological changes in the R859C channel [88]. The model used here (ModelDB accession 87585) implements two of them. First, activation shifts in the depolarizing direction, so that a stronger depolarization is needed to open the channel: the source reports an 8.5 mV shift of the activation midpoint, and the mechanism approximates the measured relationship with a +6.1 mV midpoint shift together with an approximately 34% change in slope factor, since a pure translation of the curve does not reproduce it. Second, mutant channels carried markedly smaller currents, a *∼*10-fold decrease when co-expressed with *β*1 subunits; this reduction is not represented in the model, whose maximum sodium conductance is identical in the wild-type and variant mechanisms. Third, recovery from slow inactivation was substantially slower, the slow time constant rising from 40.5 s to 141.7 s and the fraction of current recovering by the slow route from 47% to 57%; the mechanism reproduces these constants at the *−*120 mV recovery potential at which they were measured, but at the *−*60 mV resting potential of the protocol used here the two time constants are 131 s and 140 s, both far longer than the 300 ms sweep, so this difference does not act within the simulation. Figure 5 therefore shows the consequence of the activation shift alone.

**Figure 5:**
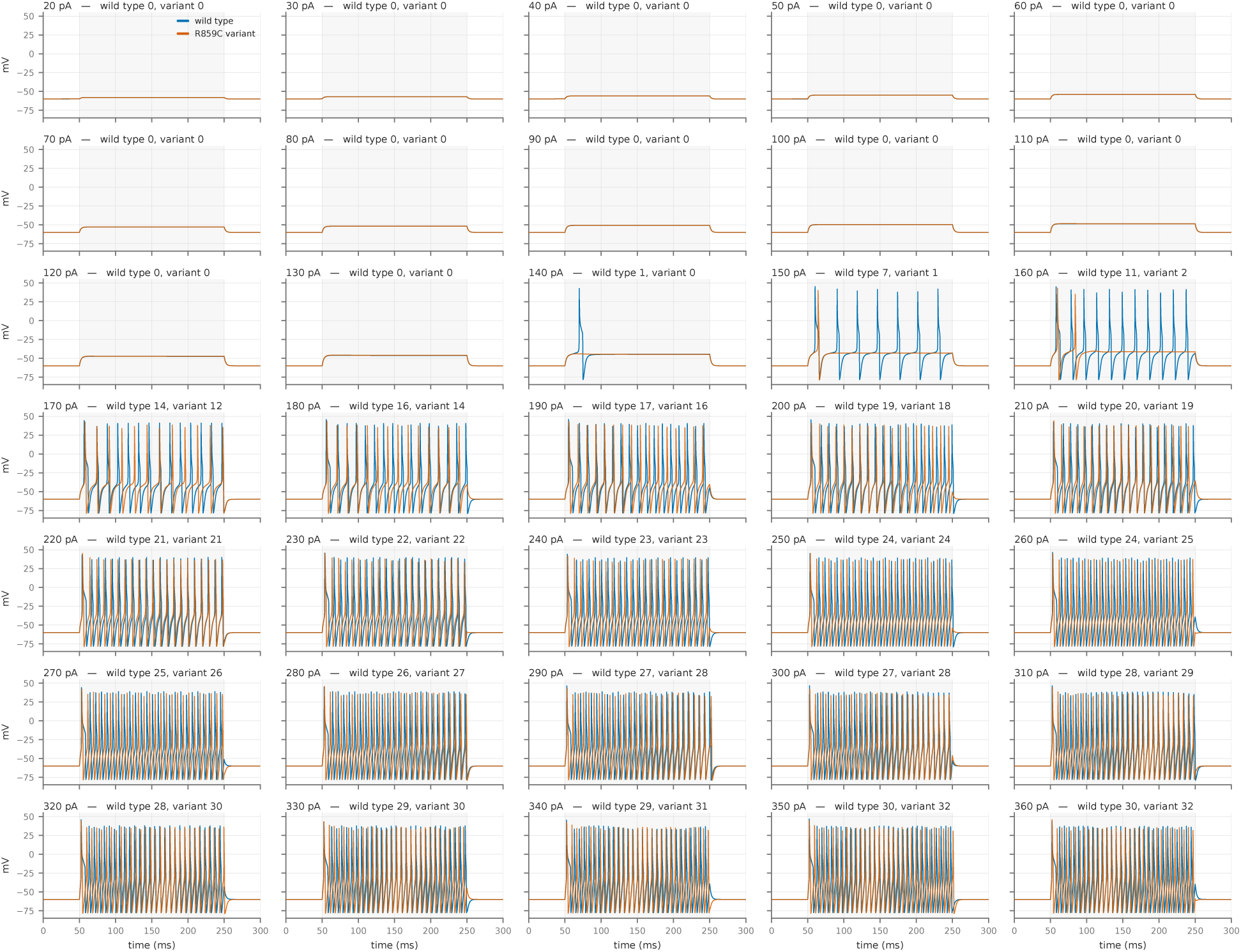
The three regions of the R859C deficit are visible trace by trace across the stimulus ladder. Thirty-five panels, one per injected-current level from 20 to 360 pA, with reference and variant overlaid, the 200 ms current step beginning at 50 ms shaded, and the spike count for both arms in each panel title. Traces are the archived NEURON output, integrated at a 0.01 ms step and logged every tenth step, giving 3,001 samples over 300 ms; the time axis is derived as sample index multiplied by the 0.1 ms logging interval rather than assumed, and is labelled in milliseconds, with voltage in mV and the injected current of each panel in pA. Both cells are silent from 20 to 130 pA. Separation appears between 140 and 210 pA, the two arms coincide from 220 to 250 pA, and the variant fires the additional action potentials above 260 pA.

Computational modeling incorporating these experimentally determined channel properties (Figure 5) revealed that these alterations collectively increase neuronal firing thresholds and reduce repetitive firing capacity. Specifically, at 150 pA the wild-type cell fires 7 action potentials and the variant fires 1; the variant reaches 2 only at 160 pA. This reduction follows from the shift in activation. Reverting only the activation midpoint and slope to their wild-type values, and leaving the variant’s slow-inactivation kinetics in place, restores the wild-type spike count at all thirty-five levels; substituting the wild-type slow-inactivation time constant instead alters one action potential at one level, because that process has a time constant of 131 to 140 s at the resting potential used here against a 300 ms sweep.

Both cells are silent from 20 to 130 pA. Measured across the full 20–360 pA ladder, however, the deficit is regional rather than global, and falls into three contiguous regions: the variant is less excitable from 140 to 210 pA, indistinguishable from the reference at every level from 220 to 250 pA, and more excitable at all eleven consecutive levels from 260 to 360 pA. Summed over the ladder the two cells fire almost the same number of action potentials – 498 for the wild type against 490 for the variant. The R859C variant therefore shifts the threshold for repetitive firing rather than abolishing the capacity for it, which is a different therapeutic target from a global loss of excitability.

### 3.2 A Kv7.2 Variant Reproduces a Published Hyperexcitability Phenotype and Is Compensated Without Touching the Mutated Channel

To test whether the same framework transfers to a different channel family on an independently published model, we built the potassium case study on the multi-compartment CA1 pyramidal neuron model of Miceli et al. [89] (ModelDB accession 118986), which supplies the M-current mechanism in wild-type and Kv7.2 D212G variant forms. The variant causes benign familial neonatal seizures (BFNS), a condition in which loss of the M-current removes a brake on repetitive firing [91–93]. Loss-of-function in these subunits arises through more than one molecular route: altered channel gating that shifts the activation curve [91], and reduced current when mutant and wild-type subunits co-assemble [94]. The magnitude of the resulting current change correlates with long-term neurodevelopmental outcome [94, 95].

We began by reproducing the figure distributed with the model before building anything on top of it. Under the model’s own published protocol (a 0.47 nA somatic step from 5 to 405 ms, 500 ms total, at 35 °C), the reference cell fires a single action potential, at 11.9 ms, while the variant fires eight, at 10.8, 46.4, 98.8, 152.4, 206.5, 260.1, 313.8 and 367.7 ms. Both the count and the individual spike times match the screenshot shipped inside the model package (Figure 6). The reproduction establishes that the variant phenotype compensated below is the published one rather than an artefact of our own configuration.

**Figure 6:**
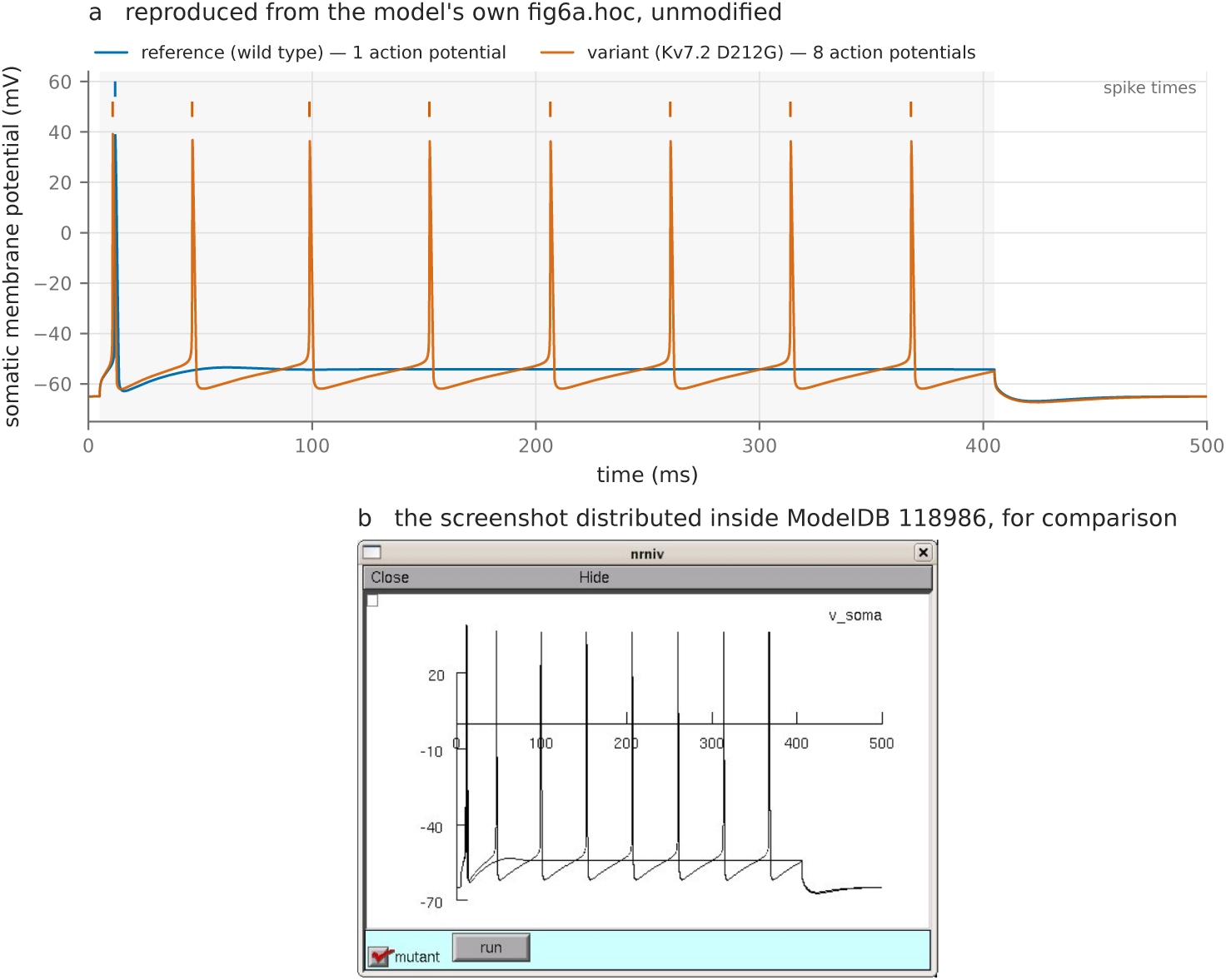
The Kv7.2 D212G phenotype reproduces the published result under the model’s own protocol. **(a)** Our simulation of the multi-compartment CA1 pyramidal model of Miceli et al. [89] (ModelDB accession 118986) under the protocol shipped with it: a 0.47 nA somatic current step from 5 to 405 ms, 500 ms total, at 35 °C, with a single flag selecting which mechanism carries the M-current. The reference cell fires once, at 11.9 ms; the variant fires eight times, at 10.8, 46.4, 98.8, 152.4, 206.5, 260.1, 313.8 and 367.7 ms. **(b)** The screenshot distributed inside the model package, shown as the verification target; it is the model author’s own file and is reproduced here for comparison only. Both the spike count and the individual spike times agree, which establishes that the phenotype compensated below is the published one.

The form of the variant matters for the therapeutic argument. The maximal conductance of the M-current is *identical* in the two arms of the model, at 0.0001 mho/cm^2^; the mechanisms differ in nine gating parameters alone: the activation midpoint and slope, the rate coefficients, and their voltage dependences. The Kv7.2 variant is therefore the same form of defect as R859C, altering channel kinetics rather than channel expression, in a different channel family and on an independent published model.

Across a ladder of twelve injected-current levels spanning 0.30 to 1.00 nA, the untreated variant fires 133 action potentials in total against the reference cell’s 97 (Figure 7). The separation is concentrated between 0.44 and 0.50 nA, where the untreated variant fires 7 to 9 action potentials at levels where the reference fires 1 or 2: the loss of the M-current removes the brake precisely in the range where the reference cell is still near threshold.

**Figure 7:**
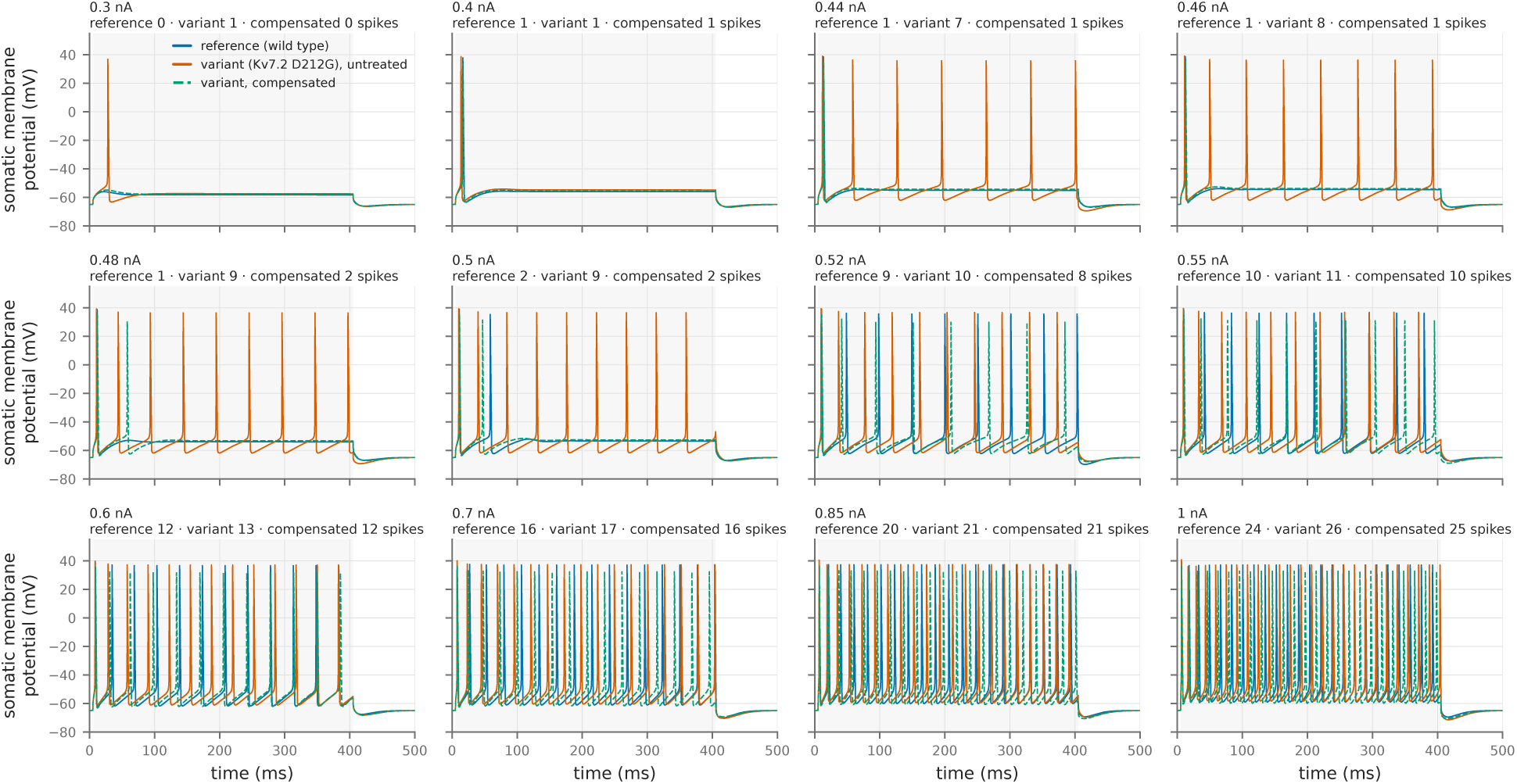
The compensated Kv7.2 cell tracks the reference at every level of the stimulus ladder. Twelve injected-current levels from 0.30 to 1.00 nA on the multi-compartment CA1 pyramidal model (ModelDB accession 118986), with the reference, untreated variant and compensated arms overlaid and per-panel spike counts for each. The current step begins at 5 ms and lasts 400 ms of a 500 ms sweep (shaded), at 35 °C; action potentials are counted by upward crossing of 0 mV at the soma. The compensated arm applies a 0.79-fold change in sodium conductance, a 1.94-fold change in delayed-rectifier potassium conductance and a 0.97-fold change in leak conductance, with the mutated M-current untouched. Totals across the ladder: reference 97 action potentials, untreated variant 133, compensated variant 99. Time is labelled in ms, voltage in mV, and each panel title carries the injected current in nA.

Adjusting only the pharmacologically accessible conductances, and leaving the mutated M-current untouched, restores 88.9% of the reference firing pattern (Figure 8). The prescription is a 0.79-fold change in sodium conductance, a 1.94-fold change in delayed-rectifier potassium conductance and a 0.97-fold change in leak conductance; the compensated cell fires 99 action potentials against the reference’s 97, and tracks the reference across the ladder. The measure is a count, so the figure is quantized: the untreated variant differs from the reference by 36 action potentials over the ladder, one action potential is worth 100/36 = 2.78 percentage points, and 88.9% is accordingly quoted to one decimal place, which is the precision the data supports. Configurations of this quality are also rare. Of 2,000 candidates drawn, 1,864 were scored (the remainder were rejected as stiff or divergent), and of those one exceeded 80% restoration, two exceeded 70% and nine exceeded 60%. Effective compensation therefore exists and is reachable by search, and reporting the full distribution alongside the best candidate is what tells a reader how much search it takes to find one.

**Figure 8:**
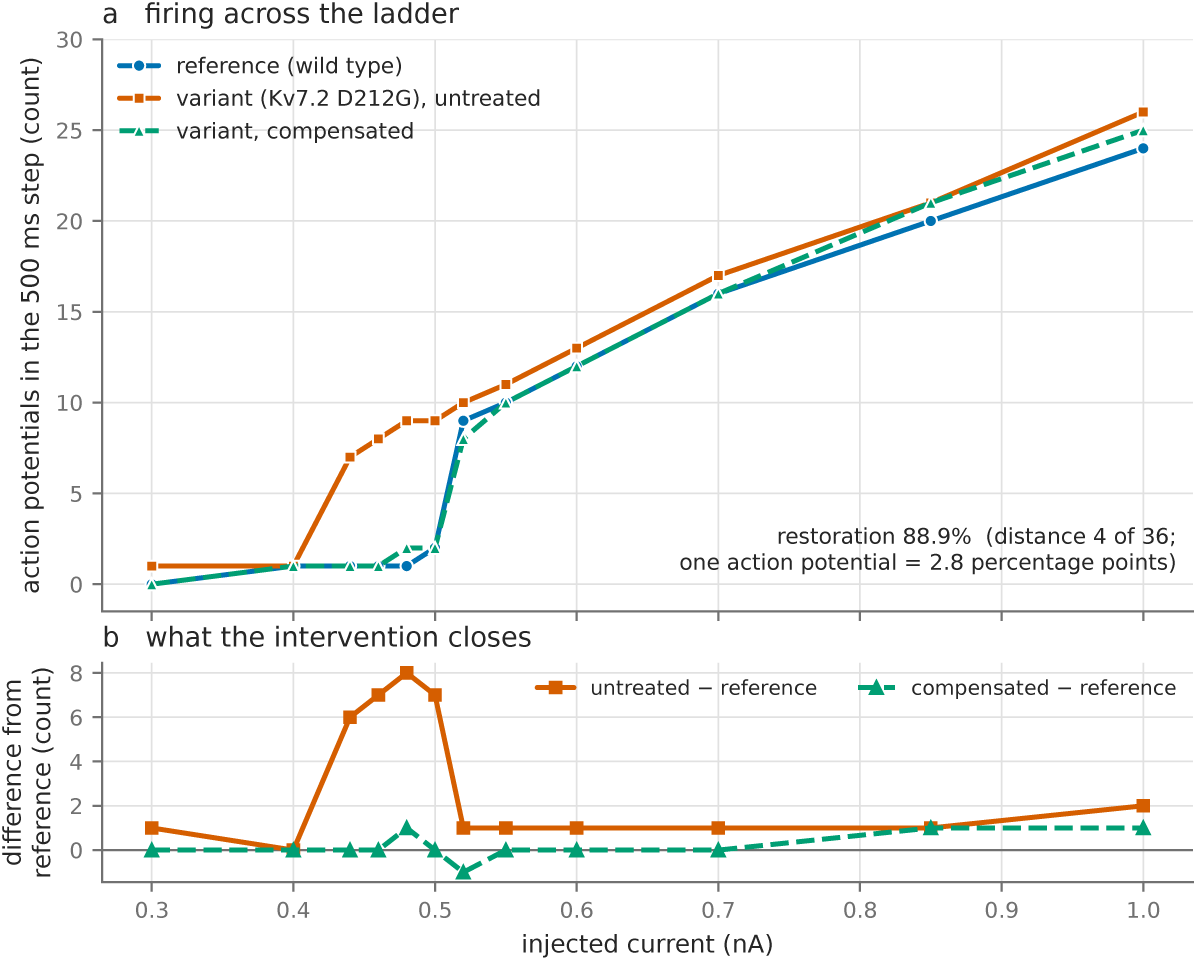
Adjusting only the pharmacologically accessible conductances restores the reference firing pattern. **(a)** Spike count against injected current for the reference, the untreated variant and the compensated variant across the twelve-level ladder. **(b)** The per-level difference from the reference for the untreated and compensated arms, which is the quantity the intervention closes. The mutated M-current lies outside the search space and its maximal conductance is identical in both arms of the model; the prescription changes sodium conductance by 0.79-fold, delayed-rectifier potassium conductance by 1.94-fold and leak conductance by 0.97-fold, and recovers 88.9% of the reference pattern. One action potential is worth 2.78 percentage points on this measure, which is the precision the data supports. Candidates were drawn log-uniformly; 1,864 of 2,000 were scored, the remaining 136 being rejected as stiff or divergent.

### 3.3 Compensating Configurations Occupy an Extended Region of Conductance Space

We explored the parameter space, examining the relationship between conductance modifications and the restoration of neuronal behaviour.

We mapped the solution topology on the differentiable model, whose throughput is what makes a survey of this size affordable: a lattice of 8,000,000 conductance triples per variant, evaluated for four variant types (the R859C sodium-channel shift, the Kv7.2 D212G potassium-channel shift, a larger potassium shift, and both shifts applied together), decomposed at 33 to 34 selection thresholds each. The four surveys together are 32,000,000 forward evaluations across all 35 injected-current levels, and each variant took 274 seconds on a single RTX 2070; at the non-differentiable pipeline’s measured node rate the same survey would take 7.8 days per variant. Reporting the decomposition across a continuous range of thresholds separates the apparent dimensionality of the solution set from the stringency of the selection used to define it. We assessed efficacy by spike count for overall activity and by DTW distance for voltage trajectory shape.

The parameter sets that restore reference behaviour do not form isolated points. They occupy an extended, low-dimensional structure in conductance space, and we characterise it by principal-direction decomposition at a continuous range of selection thresholds (Figure 9). Across every threshold examined, and for sodium-channel and potassium-channel kinetic variants alike, the third principal direction accounts for less than 12% of the variation when the decomposition is performed on the logarithm of the conductances, which is the space in which the parameter box is defined and sampled, and for under 5.5% at any selection retaining less than one percent of the spiking candidates, under 3.5% for the two clinically modelled variants. In linear coordinates the same statistic reaches 20.4%. Selection thresholds are quoted throughout as the share of spiking candidates retained by an inclusive spike-count distance cut. The set never fills a volume at any threshold examined. The number of candidates retained ranges from 6 to 360,317 across the thresholds swept; at the five tightest selections the decomposition is fitted to between 6 and 31 points, and those rows should be read as showing that the selected set is flat rather than as measuring how flat. A threshold on a scalar criterion over three parameters yields a selected set that is generically close to two-dimensional whatever the criterion is: across 24 smooth criteria with no compensation structure, evaluated on the same lattice at the same selection sizes for all four variant types, the third principal direction has a median between 0.08% and 18.6%, and the compensating set’s own value sits inside that range for every variant. We therefore report the extent of the compensating set, which bears on whether an intervention can reach it, and not its dimensionality.

**Figure 9:**
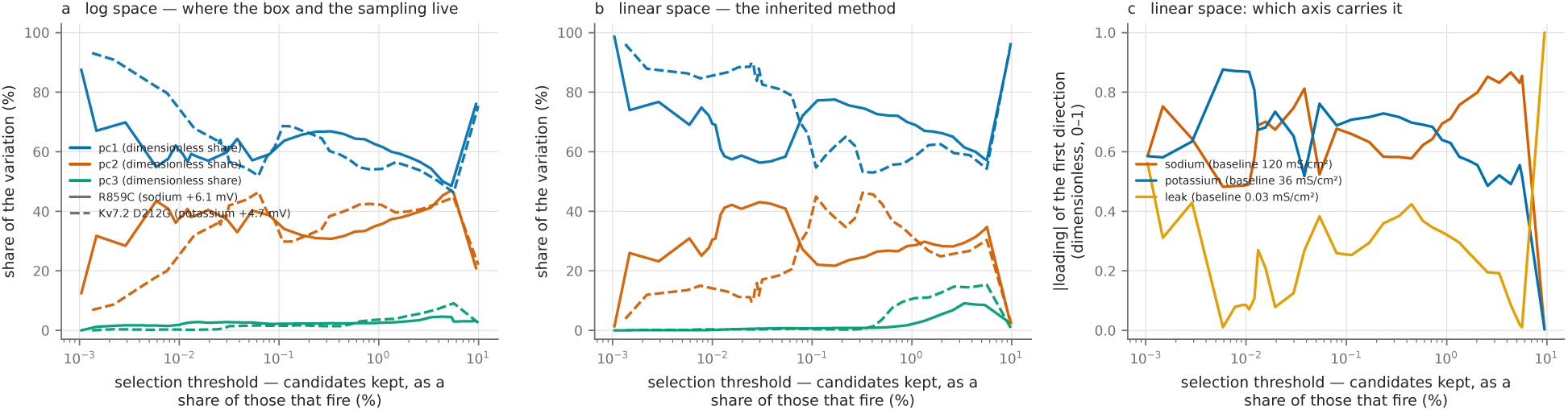
Compensating configurations occupy an extended region at every selection threshold. Principal-direction decomposition of the selected set against the selection threshold, for four variant types, each on a lattice of 200^3^ = 8,000,000 conductance triples evaluated at all 35 injected-current levels. Selection retains every candidate within a given spike-count distance of the reference rather than a fixed number of best candidates, because the score is an integer and candidates arrive in large tie blocks; thresholds are quoted as the share of spiking candidates retained. Each axis is normalised by its own range before the decomposition. **(a)** Decomposition on the logarithm of the conductances, the space in which the box is defined and sampled. The third principal direction stays below 12% at every threshold and below 5.5% at any selection retaining less than one percent of the spiking candidates, and below 3.5% for the two clinically modelled variants. What varies with the threshold is whether one direction dominates or two do; the four variant types are indistinguishable at matched thresholds. **(b)** The same decomposition in linear coordinates, where the second direction collapses at loose selections, an artefact of the coordinate system that appears for no variant in log coordinates. **(c)** The conductance axis carrying the linear first direction, which moves with the threshold and at the loosest selections lies on the leak axis, the axis with the narrowest range in decades.

What changes with the selection threshold is how the extent is distributed among the directions: at the tightest selection examined, which retains the top 0.01% of spiking candidates, a single direction carries 60–85% of the variation, and as the threshold is relaxed the structure opens smoothly into two comparable directions, approximately 48% and 45% at the top five percent (Figure 9). The four variant types tested are indistinguishable on this measure at matched thresholds, differing by 8–10 percentage points with no consistent ordering by channel family; an apparent contrast between mutation types is therefore a property of the threshold applied rather than of the mutation. The structure supports a claim about accessibility rather than about geometry: compensating configurations are not isolated points, so an intervention need not reproduce a single parameter combination exactly, and the apparent shape of the set follows from how tightly it is selected rather than from the mutation.

This threshold-dependent structure means multiple parameter combinations produce functionally equivalent outcomes. This phenomenon aligns with recent experimental demonstrations using dynamic clamp techniques, which revealed that diverse Hodgkin-Huxley parameter sets can map to identical functional coordinates in reduced-dimensional phase space, maintaining equivalent excitability properties despite vastly different underlying molecular configurations [96, 97]. This carries a direct therapeutic implication: a successful treatment need not restore the original parameter values, but can guide the system toward any of the functionally equivalent states within the compensating set.

### 3.4 R859C MT-MT Enumeration Defines Alternative Ground-Truth Configurations

To determine whether the baseline conductance configuration is a unique solution or one of several states that reproduce the same neuronal behaviour, we enumerated the MT-MT parameter space for the R859C variant.

Thirty-seven distinct conductance configurations lie at zero distance from the observed variant behaviour under the spike-count measure, read directly from the stored candidate database. Each was carried through the full intervention analysis, so that every candidate intervention is tested against all 37 rather than against the experimentally matched baseline alone.

These 37 triples are alternative candidate ground truths for the variant cell rather than alternative therapies: the observed R859C behaviour does not identify a unique conductance triple under this measure, and the experimentally matched baseline is one admissible configuration among the 37. Enumerating them is what allows a candidate intervention to be required to work across all of them, so that the analysis tests compensation across a set of admissible ground-truth models rather than against a single assumed molecular configuration.

Thirty-seven conductance triples fire the same number of action potentials at every one of the 35 injected-current levels. That is degeneracy, and it is what renders the therapeutic target tractable. Exact restoration of a single parameter set is a demanding requirement for drug development; because equivalent spike-count behaviour arises from more than one conductance configuration, a candidate intervention can be required to succeed across all 37 admissible ground truths and still be identified. The enumeration fixes which configurations are admissible, and the intervention analysis scores every candidate against the full set.

### 3.5 The Efficacy Distribution Runs Smoothly to a Best of 94.6% With No Functional Ceiling

We evaluated every candidate intervention constructed for the R859C variant and examined the resulting distribution of outcomes.

Each of the 1,050 MT-WT candidates was paired with each of the 37 MT-MT configurations, and each resulting difference vector applied to each configuration, giving 1,437,450 intervention tests actually simulated under the spike-count criterion. We report tests simulated rather than a doubled upper bound: both an additive and a subtractive direction were attempted, but under this criterion the subtractive direction contributed no surviving tests at all, for a structural reason: every difference vector is positive in all three conductances while the 37 variant configurations are smaller, so every subtraction produces a negative conductance and is rejected before simulation.

Of the simulated tests, 1,375,599 (95.70%) score above the *−*100% floor of Equation (4), meaning they leave the cell less than twice as far from the reference as no treatment, and are plotted; the remaining 61,851 fall below the floor and are discarded. The dynamic time warping criterion was applied to its own candidate set of 70 MT-WT candidates and 70 MT-MT configurations, giving 4,900 difference vectors. Of these, 1,606 are positive in all three conductances, so both directions survive in part: of the 686,000 candidate combinations, 560,230 have no negative conductance and were simulated. The efficacy results in this section are on the spike-count criterion; the time-warping conditions supplement it and their distribution is not reported here.

The distribution (Figure 10) bounds what this intervention set achieves. The best intervention reaches 94.59% similarity (a residual distance of 2 action potentials out of 37, with 10,780 tests tied at that value), and no intervention anywhere in the set reaches 100%, or even 97%. Those 10,780 tests are 10,780 distinct conductance triples, but they produce only six distinct actionpotential-count profiles across the 35 stimulus levels, and 9,102 of them (84.4%) share a single profile that spans the full seven-fold range of prescribed potassium conductance present in the set. The redundancy therefore extends to behaviour and not only to score. The dominant profile matches the reference ladder’s action-potential count exactly at 33 of the 35 stimulus levels, and fires one additional action potential at each of the remaining two, at 260 and 290 pA. No intervention in this set achieves complete replication of reference behaviour, and that bound now rests on the full 1,437,450 tests.

**Figure 10:**
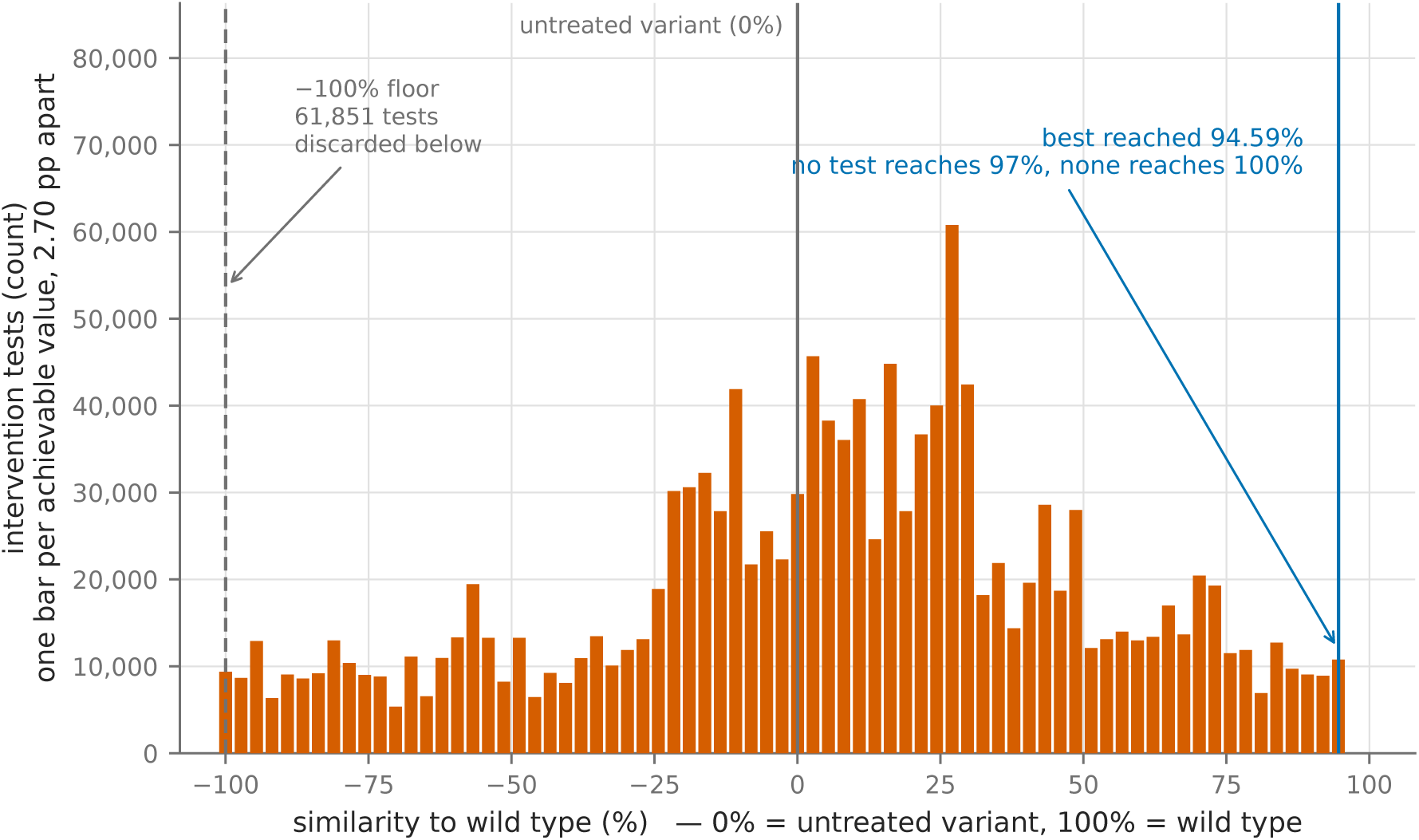
Efficacy is quantized by the action-potential count, and the best intervention stops two steps short of full restoration. All 1,437,450 tests simulated for the R859C variant under the spike-count criterion: 1,050 MT-WT candidates paired with 37 MT-MT configurations give 38,850 interventions, each applied to each of the 37 configurations. Similarity is computed as in Equation (4), with the untreated variant at 0%. Bars are placed on the measure’s own quantum, since the spike-count distance is an integer number of misplaced action potentials and similarity can therefore take only values 100/37 = 2.70 percentage points apart. 1,375,599 tests (95.70%) clear the *−*100% floor and are plotted; the remaining 61,851 fall below it. The best value reached is 94.59%, no intervention reaches 97%, and 165,252 tests (11.50%) exceed 60% with no inflection there.

The distribution is continuous over its whole range, with no threshold beyond which efficacy stops improving: 165,252 interventions (11.50%) exceed 60% similarity and 38,474 (2.68%) exceed 85%, and the density falls away smoothly towards the maximum with no inflection anywhere along the scale. The median plotted intervention reaches 8.11%, with the tenth and ninetieth percentiles at *−*62.16% and 64.86%. What limits efficacy here is the thinness of the upper tail rather than a ceiling, and the therapeutic consequence is favourable: the number of high-efficacy candidates recovered grows with the number of candidates evaluated, so enlarging the search, which the differentiable forward model of Section 3.7 makes affordable, is a productive route to a better prescription.

Three references appear in this work and each carries its own scale. Efficacy divides by 37, the separation between the variant and the reference the search integrated alongside it, which is the reference every stored candidate distance was measured against: across the 35 injected-current levels that reference fires 497 action potentials against the variant’s 488, and their summed absolute per-level difference is 37. The archived reference ladders separate by 38 under second-order integration, and that is the figure the sensitivity comparison of Section 3.6 uses. Topology divides by 1,281, the worst score in the candidate table. The same best candidate reads 94.59% on the first scale and 99.84% on the third, and on the third a 97% cut admits every candidate in the table while a 99.5% cut still admits 70% of it, so selection thresholds on that scale do not identify high-performing candidates. A percentage from this pipeline carries meaning only when its reference is named, and we name all three.

### 3.6 The Compensating Configuration Tolerates a Median Dosing Error of *±*11% and *±*3% in the Least Favourable Direction

We tested how robust each intervention remained when conductance parameters varied from their optimal values. Perturbations are applied as a fraction of each optimized conductance rather than as an absolute increment, and only to the pharmacologically accessible conductances; the mutated channel is not being dosed and is held fixed. We report three modes: one conductance perturbed at a time, joint independent draws on all of them, and the exhaustive set of sign corners, which gives the worst combination of perturbation directions. In each case we record the largest perturbation at which efficacy remains above 80% of its unperturbed value.

On the multi-compartment model, efficacy survives a joint fractional perturbation of *±*11% at the median of the draws, *±*5% at their fifth percentile, *±*4% with one conductance perturbed at a time, and *±*3% under the worst combination of directions (Figure 11). Every integer magnitude from 1 to 20% was measured, so each window is resolved to 1%. The window measures the same property continuously: efficacy above the criterion is retained across a range of settings of all three accessible conductances, so a dose has a region to land in. The median figure is the one that describes a typical patient and the worst-case figure is the one any guarantee would have to rest on, and they differ by more than a factor of three. These curves are themselves quantized. Efficacy here is normalized to its value at the optimum rather than to the untreated variant, so one action potential is worth 3.1 percentage points on this scale, against 2.78 points on the variant-normalized scale of Section 3.2. Each crossing should be read as falling within one such step of the value given.

**Figure 11:**
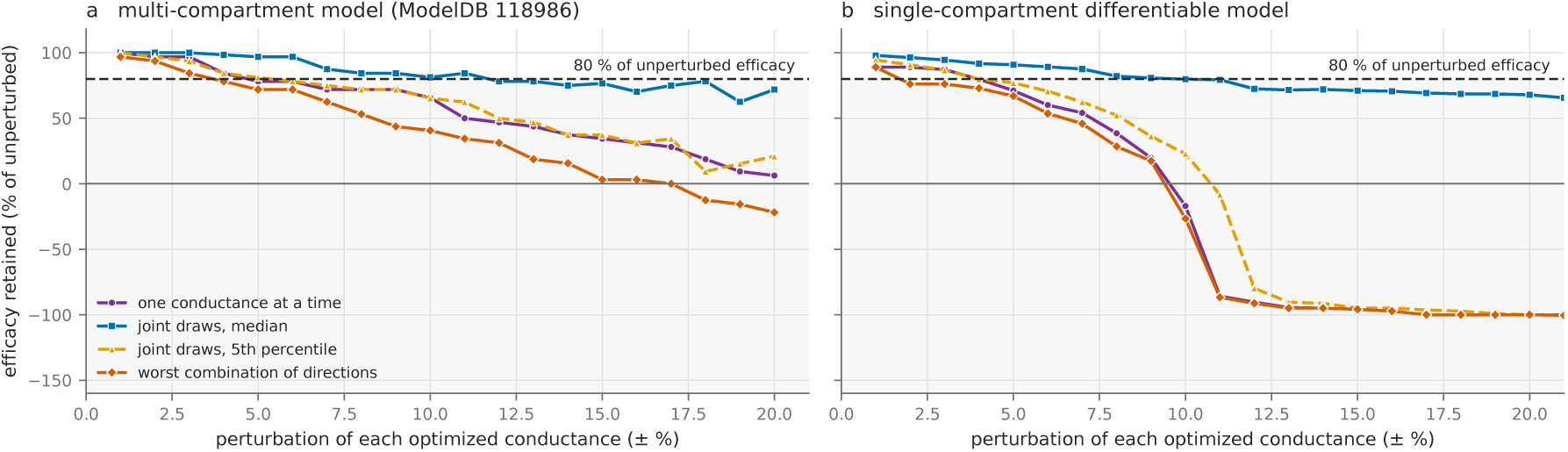
The compensating configuration tolerates a median dosing error of *±*11%, and *±*3% in the least favourable direction. Efficacy retained against perturbation magnitude, expressed as a fraction of each optimized conductance and applied only to the pharmacologically accessible conductances, never to the mutated channel, which is not being dosed. **(a)** The Kv7.2 multi-compartment model (ModelDB accession 118986), at 88.9% efficacy at the optimum, with 120 joint draws per magnitude and the one-at-a-time and sign-corner modes enumerated exhaustively. **(b)** The single-compartment differentiable comparator, the potassium variant with *ḡ*_K_ held at 20. The dashed rule marks the 80% efficacy criterion and the solid rule marks parity with no treatment; efficacy is normalised to its value at the optimum rather than to the untreated variant, so a curve below the solid rule describes a mis-dosed intervention that leaves the cell further from the reference than no intervention at all. Magnitudes are sampled on a 1% grid: every integer from 1 to 20% in panel (a) and from 1 to 50% in panel (b). Curves are quantized: one action potential is worth about three efficacy points on the multi-compartment ladder.

The single-compartment differentiable model is a conservative proxy for dosing tolerance in every mode, by a factor between 1.2 and 3. On the 1% grid now shared by both models, the windows of the Kv7.2 multi-compartment model (ModelDB accession 118986) are the wider ones in every mode: *±*11% against *±*9% at the median, *±*5% against *±*3% at the fifth percentile, *±*4% against *±*3% with one conductance at a time, and *±*3% against *±*1% under the worst combination of directions. The single-compartment model is markedly more sensitive to the same kinetic shift than the multi-compartment models, which matters wherever it is used as a proxy for them, and that greater sensitivity is a separate quantity from the ratio of the stability windows. Summed over the injected-current ladder, the absolute per-level difference in spike count between reference and variant is 459 in the single-compartment model, against a reference total of 470; in the R859C multi-compartment model (ModelDB accession 87585) the same summed difference is 38, against a reference total of 498. Neither figure is a count of action potentials lost. Each is the summed absolute difference in spike count between reference and variant across all thirty-five levels, which is the denominator *d*_variant_ of Equation (4). The multi-compartment pair is measured on the archived reference ladders under second-order integration, which is the second of the three scales set out in Section 3.5; the efficacy denominator of Equation (4) is measured against the reference the search itself integrated and is 37. In the multi-compartment model the net change is a deficit of only 8 spikes, and the variant in fact fires more at eleven of the thirty-five levels.

The measured windows place the compensating configuration in a stable region of parameter space rather than at a brittle point solution. This tolerance is what an intervention needs in order to survive biological variation, and testing it in patient-derived cells with diverse genetic backgrounds is the next experimental step.

### 3.7 A Differentiable Forward Model Makes Large Non-Local Searches Affordable

The exhaustive search described above is limited by the cost of the forward simulation, not by the difficulty of the optimization. We therefore reimplemented the Hodgkin-Huxley model in PyTorch [80, 81] rather than in NEURON [79], which makes the model both differentiable and evaluable in batch on a graphics processor. We report two findings: the throughput the fast forward model delivers, and how gradient descent compares with direct search on that same model.

#### Throughput

We define one candidate parameter set as one conductance triple simulated across all 35 injected-current levels, 300 ms each, at a 0.01 ms integration step, and report every rate on that unit (Table 1). Four comparisons, each made on unchanged hardware or on a stated change of hardware, separate the sources of the gain. On one processor core, the differentiable model evaluates 0.089 sets per second one at a time and 20.8 when 1,024 sets are evaluated together, a factor of 234 from batching alone on unchanged hardware; at that batch size it is 20.8 times NEURON’s 1.0 sets per second on the same core. Moving the batched model to an RTX 2070 graphics card reaches 543 sets per second. Compiling the integration step into fused kernels raises the same card to 34,279, a factor of 63 on unchanged hardware. A 2023 RTX 4070 Ti reaches 63,649. In production the non-differentiable pipeline sustained 11.9 sets per second on a 72-core node during the spike-count stage and 16.9 during the time-warping stage; these are end-to-end wall-clock rates for a complete production stage, which also computes the summary statistics for every candidate, and against them the fused differentiable model is between three and four orders of magnitude faster. The single-core figure of 1.0 set per second was measured on a simpler per-candidate loop in a separate job; 72 cores at that rate would give 72 sets per second against the 11.9 measured, and we report the node rate as the production figure.

Fusion speeds the loop up by moving less data, not by doing arithmetic faster. Before fusion, each of 76 kernels per integration step reads its operands from device memory and writes its result back, demanding 412–420 GB/s against the 393.6 GB/s the card can supply, so the loop runs at 105–107% of available bandwidth and stalls. Fusion removes the intermediate traffic rather than the kernel launches, and afterwards the same loop demands only 51–64 GB/s, 13–16% of measured bandwidth.

Across the two accelerator generations tested, delivered throughput rose by 1.86*×* against a 5.37*×* difference in rated arithmetic throughput. Rated performance does not predict throughput for this workload, and we therefore report measured rates only.

Four qualifications bound the comparison. The non-differentiable rate includes summary statistics that the differentiable rate does not, so the comparison favours the differentiable pathway. And the differentiable model run unbatched on a single core evaluates 0.089 sets per second, about eleven times *slower* than NEURON on the same core. The gain is therefore a property of batched evaluation on a parallel accelerator rather than of the formulation itself. Two further qualifications bound it. Hardware is not normalised: the comparison sets consumer graphics cards from 2018 and 2023 against server processor cores from 2020. And the biophysics is not identical: the nondifferentiable production mechanism carries four state variables against three in the differentiable model. This second qualification applies to the node figures; the single-core comparison is made on the same three-state model on both sides. The seven graphics-card rates in Table 1 are medians of five independent end-to-end launches of the same benchmark on the hardware named, and the table gives the full range across the five beside each; at the batch sizes quoted the range is below 0.6% of the median for every row except the cache-resident one, where it is 6.9%. The five processor-core rates are single measurements.

#### Gradient descent against direct search

Differentiability also permits gradient descent directly on the conductances, and we tested whether it recovers the compensating configuration. A gradient-descent run is counted as having found the compensating configuration when the similarity it reaches, on the scale of Equation (4), is within 5 percentage points of the best value reached by direct search over the same parameter box. This is the same criterion used for the sampling-scheme comparison in Figure 2. Under the voltage mean-squared-error objective, with the variant channel held fixed, gradient descent found the compensating configuration in one of four variant configurations. Direct search over the same parameter box found it in four of four. Expressed on the similarity scale of Equation (4): with sodium fixed, gradient descent reached 94.23% against direct search’s 96.15%; with potassium fixed, *−*87.50% against 93.97%; with leak fixed, *−*123.38% against 95.02%; and for the kinetic variant, 1.74% against 97.60% (Figure 12). Two of the four gradient results are worse than administering no treatment at all. The three conductance variants differ in severity: sodium is held at 1.17 times its reference value and potassium at 0.56 times, while leak is held at ten times its reference value. Direct search reached 95.02% on the leak variant at that same held value, so the descent results are failures of the optimizer rather than of the configuration. Widening the test to eight objectives across seven variant configurations and two search budgets did not rescue the method: the best objective succeeded in two of seven.

**Figure 12:**
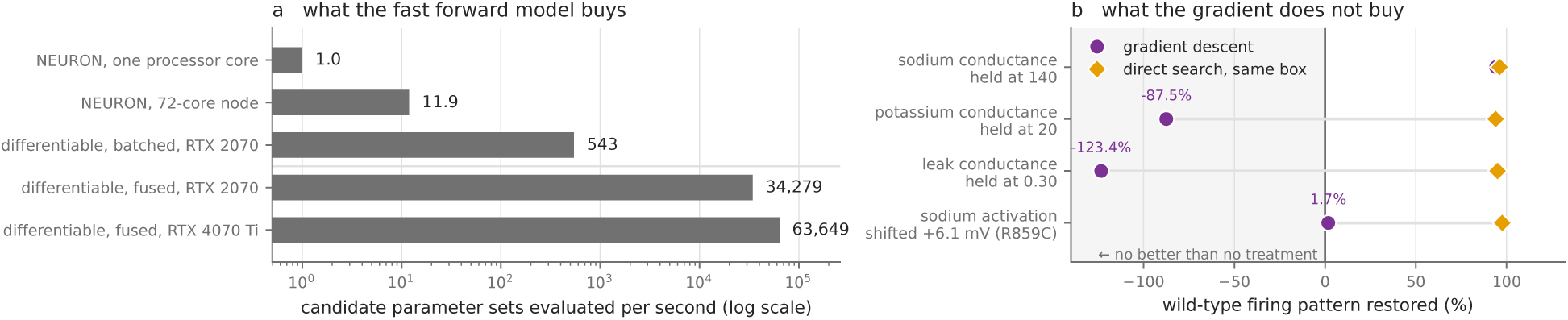
The differentiable model earns its place through forward-model throughput rather than through descent. **(a)** Candidate parameter sets evaluated per second, logarithmic axis, one bar per configuration. One candidate parameter set is one conductance triple simulated across 35 injected-current levels, 300 ms each, at a 0.01 ms integration step. The non-differentiable rate includes summary statistics that the differentiable rate does not, so the comparison favours the differentiable pathway, and no scaling projection should be drawn from these bars. Each of the three graphics-card rates is the median of five independent end-to-end launches of the same benchmark on the hardware named, as in Table 1; the two processor-core rates are single measurements. **(b)** Gradient descent under the voltage mean-squared-error loss against direct search over the same parameter box, one row per variant configuration, scored on spike count across all 35 levels and reported on the similarity scale of Equation (4). The heavy rule at 0% marks parity with no treatment. Descent reaches the compensating configuration in one case of four and prescribes a configuration worse than no treatment in two; direct search reaches it in four of four.

**Figure 13:**
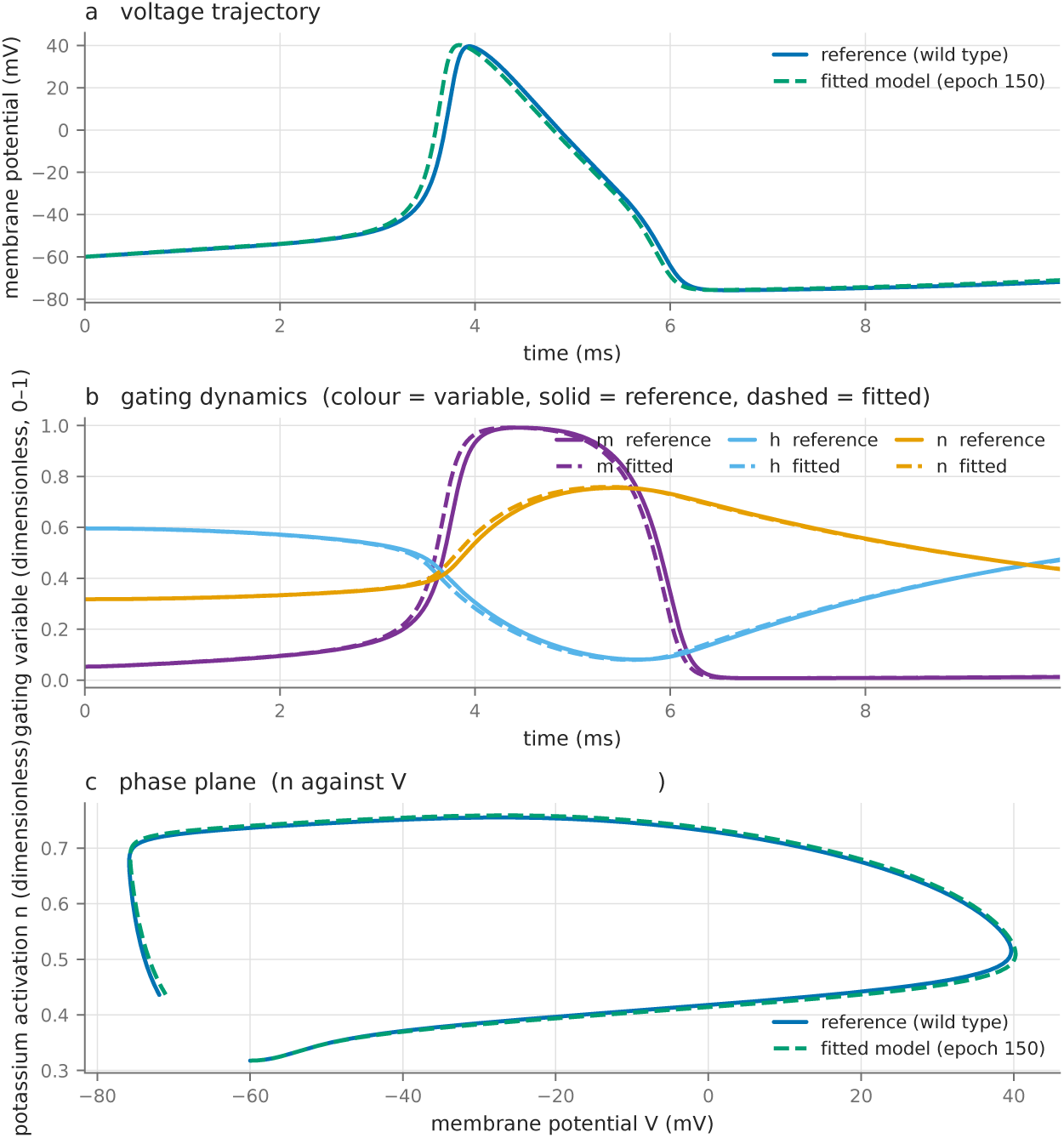
Gradient descent moves the potassium conductance onto the reference trajectory; the other two do not converge. **(a)** Membrane potential for the reference cell and the fitted model. **(b)** Gating variables *m*, *h* and *n*, one colour per variable, solid for the reference and dashed for the fitted model. **(c)** Phase plane with the potassium activation variable *n* on the ordinate against membrane potential *V* on the abscissa. Integration step 0.0066667 ms over 1,500 steps, spanning 10.00 ms, at an injected current of 8 *µ*A/cm^2^; the time vector is derived as step index multiplied by the integration step. Reference conductances (120, 36, 0.03) mS/cm^2^. The sodium conductance is held at the variant value of 140 throughout and is not fitted; potassium moves from 20 to 39.678 and leak from 0.300 to 0.155 over 150 epochs, and the epoch-150 vector is the one drawn here. The leak component is not determined by this objective and is not a converged value; see the gradient-descent results. The panel spans 10 ms and contains a single action potential.

The obstacle lies in the objective rather than in the implementation. The objective that discriminates well (spike count as a function of injected current) is not differentiable, because a spike count is an integer that does not vary smoothly with the parameters. The objectives that are differentiable are computed on the voltage trace, and their landscape is rugged enough over this parameter box that local descent reaches a poor stationary point. Differentiable relaxations of the spike count and of DTW exist (soft-DTW [98] being the best known) and are a promising route to closing this gap; among the objectives we tested, none allowed descent to succeed reliably.

For the one configuration where descent did converge, it moved the potassium conductance from 20 to 39.678 mS/cm^2^ with sodium held at the variant value, in approximately 150 iterations (Figure 13). The objective constrains the potassium conductance while leaving the leak conductance unidentified. Seven of the eight optimizers tested reached convergence, and across those seven the compensating potassium value agrees to within 2.3% of its mean, whereas the leak value spans 44.3%. We therefore report the potassium conductance as a determined result and the leak conductance solely as the epoch-150 parameter vector, not as a converged value. The remaining optimizer, Adam with AMSGrad, closed 33.0% of the loss gap, against 99.85% or more for the other seven. The phase-plane panel must be drawn at some leak value and uses the epoch-150 vector for that purpose alone, as its caption records. Convergence is better described by the stopping criterion than by an iteration count, since the count is strongly optimizer-dependent: resilient backpropagation reaches within 1% of its best loss in 31 iterations, where five of the other seven require more than 1,400.

#### Where the advantage lies

The contribution of the differentiable model is its forward pass, not its gradients. Gradient descent does not solve the compensation problem: on the objective this model admits, it succeeds in a minority of cases and can prescribe an intervention worse than none. The forward evaluation is between three and four orders of magnitude faster per candidate than the production pipeline, which makes a very large, non-local search affordable. That search, on the same fast model, succeeds where descent does not.

## 4 Discussion

### 4.1 A Systematic Framework Enables Rational Design of Ion Channel Mutation Interventions

Although we employed two distinct computational frameworks, high-fidelity NEURON simulations for clinical cases and differentiable Hodgkin-Huxley models for rapid theoretical screening, both identified extended sets of compensating configurations rather than unique parameter solutions (Figure 14). That is the property a therapy needs: an intervention has to land in the set, not on a point. We do not claim more than this from the shape of the set. A threshold on a scalar criterion over three parameters produces a nearly two-dimensional selected set whatever the criterion is, so the low dimensionality of the compensating set is not by itself evidence of biological degeneracy.

**Figure 14:**
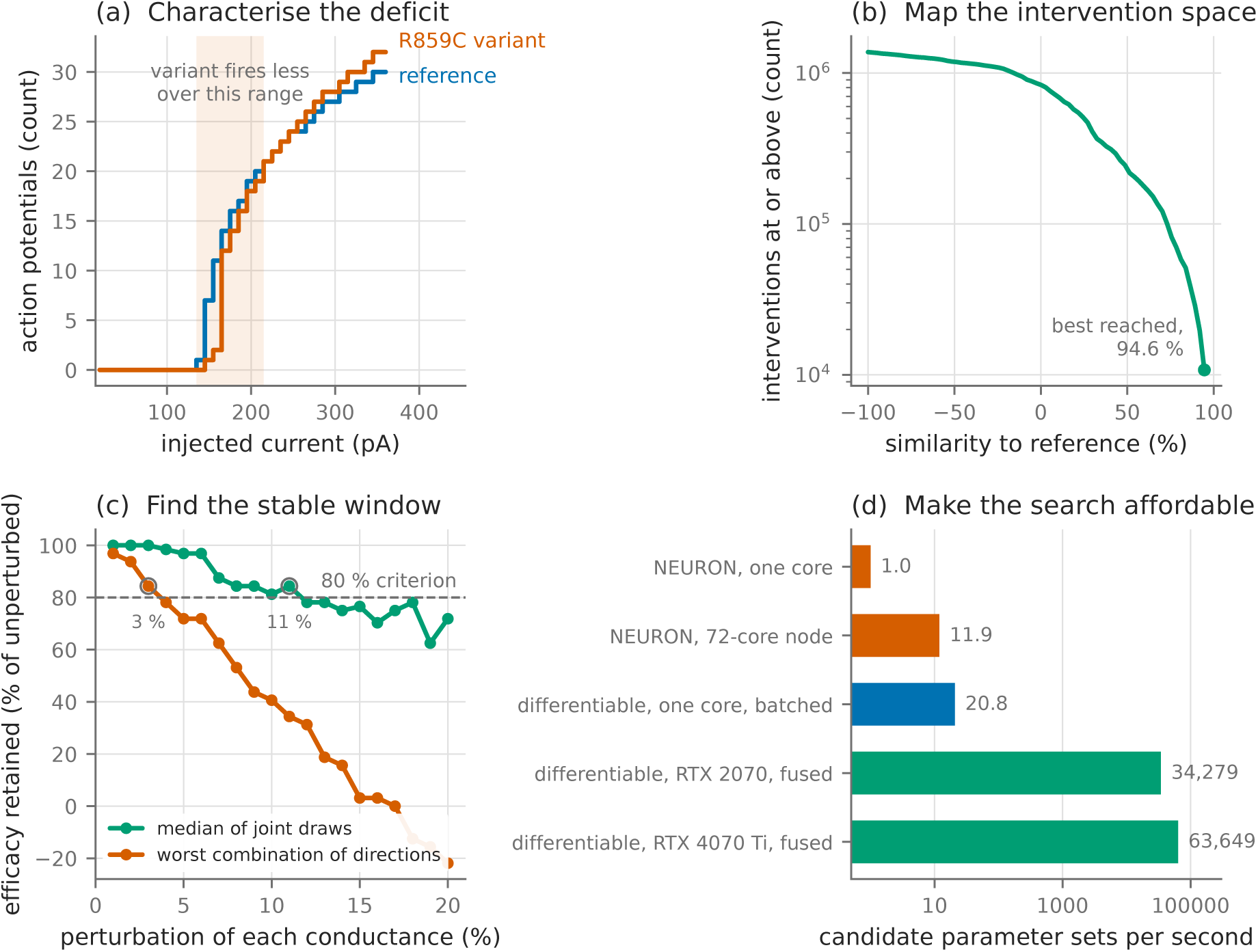
Each step of the framework produces a measured result, demonstrated across two channelopathies. The four panels recap the framework’s four steps, one measured result each. Panels (a) and (b) are the R859C case study, panel (c) the Kv7.2 case study, and panel (d) applies to both. **(a)** The Nav1.1 R859C variant (ModelDB accession 87585) against its reference across the injected-current ladder. The deficit is regional rather than global: the variant fires fewer action potentials only between 140 and 210 pA, is indistinguishable from 220 to 250 pA, and fires more from 260 to 360 pA. **(b)** Interventions reaching at least a given similarity to the reference, over the 1,375,599 tests that survive the *−*100% floor of the 1,437,450 simulated across all 37 admissible ground-truth configurations. The curve runs smoothly to its best value of 94.6% with no functional ceiling. **(c)** Efficacy retained under fractional perturbation of the pharmacologically accessible conductances in the Kv7.2 D212G multi-compartment model (ModelDB accession 118986), which reaches 88.9% efficacy at the optimum with the mutated channel untouched. Efficacy is normalised to its value at the optimum, so zero is parity with no treatment. Magnitudes are sampled on a 1% grid from 1 to 20%, and the circled points mark the largest magnitude retaining 80% of unperturbed efficacy in each mode. **(d)** Candidate parameter sets evaluated per second, logarithmic axis, on the unit defined in Section 3.7. Every graphics-card rate is the median of five independent end-to-end launches on the hardware named and every processor-core rate is a single measurement, as in Table 1. No panel reports a new measurement: each is drawn from the data file of the figure that reports it.

A critical challenge in translating computational predictions to clinical therapy is biological variability. Our topological analysis of the solution space, combined with systematic sensitivity testing, provides a data-driven assessment of robustness. The parameters that restore reference behaviour occupy an extended region of conductance space, and efficacy is maintained under fractional perturbations of *±*11% at the median of joint draws and *±*3% under the most adverse combination of directions. This indicates that therapeutic agents need not achieve pinpoint accuracy in conductance modulation; provided the intervention shifts the system into the identified stability cluster, the physiological correction is maintained.

This investigation establishes a computational framework for identifying and developing therapeutic interventions for ion channel mutations. We analysed two distinct channelopathies, the R859C sodium-channel mutation causing GEFS+ and the Kv7.2 (KCNQ2) D212G variant causing BFNS, and built a differentiable forward model fast enough to search the conductance space directly. Together these show that compensating configurations can be predicted computationally in two different channel families.

The framework proceeds in four steps: characterizing the mutation-specific electrophysiological alterations, mapping the parameter space of potential interventions, identifying the stable therapeutic windows, and searching that space at a rate which makes large non-local searches affordable. Successful interventions cluster within broad, stable parameter regions rather than requiring precise molecular targeting, which makes therapeutic development more practical.

Compensation is available from more than one direction: the parameters that restore reference behaviour form an extended region, so a therapy can be aimed at the region rather than at a point. The shape of that region follows from how tightly it is selected rather than from the biology, and the approach does not depend on its shape. The multiplicity is measured directly, in configurations that fire identically rather than in the geometry of a selected set. Taken together, the two case studies show that a compensating intervention can be identified computationally, bounded by a measured tolerance, and specified without modulating the mutated channel.

### 4.2 Limitations of this study

One translational step lies outside what this framework provides. The prescriptions it produces are changes in lumped conductances (so much more delayed-rectifier potassium current, so much less sodium current), and a lumped conductance is carried in a real neuron by several channel subtypes with distinct kinetics and distinct pharmacology. Mapping a conductance prescription onto a specific druggable subtype requires kinetic models of the candidate subtypes and a criterion for choosing among them, neither of which we address here. Without that step, the pipeline says which current must change and by how much, but not which compound will change it.

The two pathways also differ greatly in cost, which shapes how they are best combined. The NEURON simulation is high-fidelity and correspondingly expensive: on the unit defined in Section 3.7 the production pipeline evaluated 11.9 to 16.9 candidate parameter sets per second on a 72-core node, against 34,000–64,000 per second for the batched, fused differentiable model on a single graphics card. The natural division of labour would therefore be to use the differentiable model to narrow the region worth exploring in a particular case, and the NEURON model to resolve the combination of channels within it. In a biomedical application the compute time to a candidate is in any case small beside the duration and expense of deploying that candidate in patients: the full R859C search took 33.5 hours for the 1,437,450 spike-count conditions and a further 9.2 hours for the 560,230 time-warping conditions, both on a single 72-core node in the same job. Even 33.5 hours is a small cost when in vivo testing and eventual delivery to patients are likely to take years.

Finally, every result reported here is computational. The two models are independently published, and they reproduce the phenotypes described in those papers. The interventions themselves have not yet been tested in cells, and the two case studies use published models rather than recordings from an individual patient: the patient-specific route the workflow describes is the intended application rather than one exercised here.

### 4.3 Future work

These results leave one question open: how does the neuron’s own regulatory machinery navigate this parameter landscape? Previous mappings of the Hodgkin-Huxley phase space establish that the landscape is rugged, with solutions separated by regions of instability [96, 97], so a cell moving between admissible states crosses intermediate states belonging to neither. We propose that the cell’s controller reaches those states through a structure in which they are neighbours, and that the compensating set measured here is part of that structure: the 37 conductance triples that fire identically are states such a controller could move between, and the *±*11% window is the room it has to work in. An intervention enters the same set from outside, so degeneracy serves the cell’s regulation and the therapy alike. The proposal is testable on the quantities this framework already produces, by asking whether a regulatory rule that conserves the firing pattern traces a path through the compensating set.

This finding is an example of a broader class of phenomena related to cellular problem-solving, such as adaptation to mutations, drugs, and other stressors [99–101]. In all of these cases, the biology is revealing the presence of powerful methods used by living cells and tissues to infer physiological responses to accommodate an unexpected stimulus or influence. How cells search the enormous space of transcriptional and physiological states to rapidly identify adaptive responses is a key question of relevance across biomedicine, evolutionary biology, and the science of diverse intelligence [102–104].

## 5 Conclusion

This computational framework, demonstrated on two independently published models in different channel families and encompassing 1,997,680 simulated experimental conditions across two selection criteria for the R859C case study alone, provides a foundation for rational drug design and personalized medicine approaches to ion channel disorders. Future work will focus on validating these stability windows in in-vivo animal models to confirm the predicted therapeutic efficacy, and extending the model to use cases beyond neurons: the emerging control of developmental bioelectricity pathways for applications in birth defects, injury, and cancer [29, 105]. The use of computational approaches coupled with bioelectrical simulators [71, 72, 75, 76, 106], and the growing data on drugs, nanomaterials, and optical stimuli as regulators of ion channel function, represent an immense opportunity for electroceuticals targeting a wide range of biomedical and bioengineering applications [45, 46, 52, 67, 68, 107–109].

## Author Contributions

H.H.: conceptualization, methodology, software, validation, formal analysis, investigation, data curation, visualization, and writing of the original draft. M.L.: conceptualization, methodology, resources, supervision, funding acquisition, and review and editing of the manuscript. Both authors read and approved the final manuscript.

## Use of Large Language Models

H.H. used Claude (Opus 5, Anthropic) and Gemini (3.1 Pro, Google) as agentic coding assistants throughout this project: for experiment orchestration, including SLURM script generation and result intake pipelines; for data visualization; for manuscript drafting and structural revision; and for compiling technical details from the codebase into the descriptions given in Methods. These tools were not used for research ideation or experimental design. All AI-generated outputs, including code, prose and technical specifications, were reviewed, verified and refined by H.H., who takes full responsibility for the accuracy and integrity of this work.

## Acknowledgements

This work was supported by Morphoceuticals, Inc. We are deeply grateful for their generous support, which made this research possible. The authors acknowledge the use of the Tufts University High Performance Compute Cluster (https://it.tufts.edu/high-performance-computing) for the computational components of this research. Furthermore, we acknowledge Adam G. Peters, Chuhang Xiang, and Jason Xu for their help with parameter searching and basic genetic algorithms.

